# Cell type-specific inhibitory modulation of sound processing in the auditory thalamus

**DOI:** 10.1101/2024.06.29.601250

**Authors:** S Rolón-Martínez, A.J. Mendoza, C.F. Angeloni, N.W. Vogler, A.C. Drotos, M. Aizenberg, R. Chen, K. Vu, J.S. Haas, M.N. Geffen

**Author notes:** These authors contributed equally (S Rolón-Martínez, A.J. Mendoza, J.S. Haas, M.N. Geffen). Corresponding author: Maria Neimark Geffen, Ph.D. (she/her) Department of Otorhinolaryngology Perelman School of Medicine University of Pennsylvania 5 Ravdin 3400 Spruce St. Philadelphia PA 19104.

## Abstract

Inhibition plays an important role in controlling the flow and processing of auditory information throughout the central auditory pathway, yet how inhibitory circuits shape auditory processing in the medial geniculate body (MGB), the key region in the auditory thalamus, is poorly understood. The MGB gates the flow of auditory information to the auditory cortex, and it is inhibited largely by the thalamic reticular nucleus (TRN). The TRN contains two major classes of inhibitory neurons: parvalbumin (PV^TRN^)-positive and somatostatin (SST^TRN^)-positive neurons. PV and SST neurons have been shown to play differential roles in controlling sound responses in auditory cortex. In the somatosensory and visual subregions of the TRN, PV^TRN^ and SST^TRN^ neurons exhibit anatomical and functional differences. However, it remains unknown whether and how PV^TRN^ and SST^TRN^ neurons differ in their anatomical projections from the TRN to the *auditory* thalamus, and whether and how they differentially modulate activity in the MGB. We find that PV^TRN^ and SST^TRN^ neurons exhibit differential projection patterns within the auditory thalamus: PV^TRN^ neurons predominantly project to ventral MGB, whereas SST^TRN^ neurons project to the dorso-medial regions of MGB. Optogenetic inactivation of PV^TRN^ neurons bidirectionally modulated sound-evoked activity in MGB, increasing firing in 29% of MGB neurons, while suppressing firing in 41%. In contrast, inactivating SST^TRN^ neurons largely suppressed tone-evoked activity in MGB neurons. Cell type-specific computational models identified candidate circuit mechanisms for generating the differential effects of TRN inactivation on MGB sound responses. These distinct inhibitory pathways within the auditory thalamus reveal cell type-specific organization of thalamic inhibition in auditory computation.

## INTRODUCTION

Inhibition is crucial to sensory information processing and transmission in the brain. In the auditory system, inhibition plays a critical role in shaping auditory computations, from sound localization in the brainstem to statistical inference in the auditory cortex^1–7^. There are many different inhibitory neuron types within the central auditory pathway which differ morphologically, electrophysiologically, and functionally. In auditory cortex, two major sub-classes of inhibitory neurons, parvalbumin-(PV) and somatostatin-(SST) positive neurons, play distinct functional roles^8–16^, and activating PV or SST neurons leads to a decrease in frequency selectivity and an overall increase in tone responsiveness of cortical excitatory neurons^10,13,17,18^. SST, but not PV, neurons contribute to stimulus-specific adaptation, a phenomenon in which neurons adapt selectively to a subset of inputs presented frequently^8,9,19–23^.

However, cortical neurons are not the only cells that receive targeted inhibition along the auditory pathway. The medial geniculate body (MGB), located within the thalamus, receives auditory information from the auditory brainstem, and MGB neurons shape and relay sensory information to auditory cortex^24,25^. This ascending projection from MGB to auditory cortex is shaped by inhibition from the thalamic reticular nucleus (TRN), a thin sheet of inhibitory neurons that targets and envelops the thalamus^26–30^. Anatomically, the TRN receives excitatory collaterals from the MGB as well as from corticothalamic L5 and L6 neurons, and sends feedforward inhibition back to MGB^27,31–35^. Functionally, the TRN filters relevant sensory information important for perception^36–38^ and supports optimal behavioral performance in sensory attentional tasks^39–41^.

The TRN contains multiple types of inhibitory neurons, predominantly PV and SST positive neurons. Recent findings in TRN subregions dedicated to visual and somatosensory modalities have described segregated distributions of distinct inhibitory neuron subtypes that differentially target distinct thalamic nuclei and exhibit differences in intrinsic and synaptic properties^42–45^. Thus, it is plausible that as a result of both organizational and functional differences, PV^TRN^ and SST^TRN^ neurons might differentially exert inhibitory control of sound processing in the MGB. However, the specific roles of PV^TRN^ and SST^TRN^ neurons in gating the auditory thalamocortical relay have not been systematically tested.

In this study, we examined the differential effects of PV^TRN^ and SST^TRN^ neurons on thalamic sound processing. To anatomically identify how PV^TRN^ and SST^TRN^ neurons project to the sub-nuclei of the MGB, we used viral tracing techniques to express an anterograde virus encoding a flexed fluorescent reporter in the TRN of transgenic PV-Cre or SST-Cre mouse lines and traced projections to the MGB. We found that PV^TRN^ and SST^TRN^ neurons target distinct subregions of the MGB. We used viral transfection to drive a soma-targeted inhibitory opsin in the TRN to inactivate PV^TRN^ or SST^TRN^ neurons while recording neuronal activity from neurons in the MGB of awake passively listening head-fixed mice. Inactivating PV cells resulted in a mix of enhanced and suppressed MGB responses. By contrast, inactivating SST cells suppressed the vast majority of MGB responses. To identify candidate circuit mechanisms that could underlie the unexpected bidirectional modulation resulting from inactivation of SST^TRN^ and PV^TRN^ neurons, we examined MGB activity in computational models with varied synaptic connectivity within TRN and from TRN to MGB. These simulations revealed several variations of underappreciated thalamic circuitry that can explain the experimental results, including disynaptic inhibition and divergent inhibition. Together, our results demonstrate cell and circuit specificity of inhibition within the auditory thalamus that is likely to influence auditory perception.

## RESULTS

### PV and SST neurons of the TRN project to MGB, targeting different subregions

The MGB is anatomically subdivided into three main subnuclei: the ventral division (vMGB), the medial division (mMGB), and the dorsal division (dMGB). vMGB receives lemniscal projections from the brainstem and provides the most direct auditory input to auditory cortex (AC), whereas dorso-medial geniculate regions provide higher-order information. We first imaged the distribution of PV^TRN^ and SST^TRN^ neuronal projections in MGB using viral neuroanatomical tracing methods. We injected anterograde adeno-associated viruses into the TRN of PV-Cre or SST-Cre mice and traced the axonal projections to the MGB (**Figure 1A & 1C**). We took advantage of the finding that calretinin is solely expressed in the higher-order MGB^46^ to characterize the projection patterns of both neuronal subtypes within the MGB. We labeled cell bodies in dorso-medial MGB using a calretinin antibody. Whereas PV^TRN^ neurons targeted the central portion of vMGB, SST^TRN^ neurons avoided this region. PV^TRN^ projections were stronger in the central area of ventral MGB rather than peripheral and dorso-medial regions (**Figure 1B**; *N* = 14 ROIs, p < 0.001, signed-rank test). By contrast, projections of SST^TRN^ targeted the dorsal and medial MGB (**Figure 1D**; *N* = 8 ROIs, *p* = 0.023, signed-rank test). Combined, these data demonstrate that PV^TRN^ and SST^TRN^ neurons exhibit differential projection patterns to the MGB.

**Figure 1.**
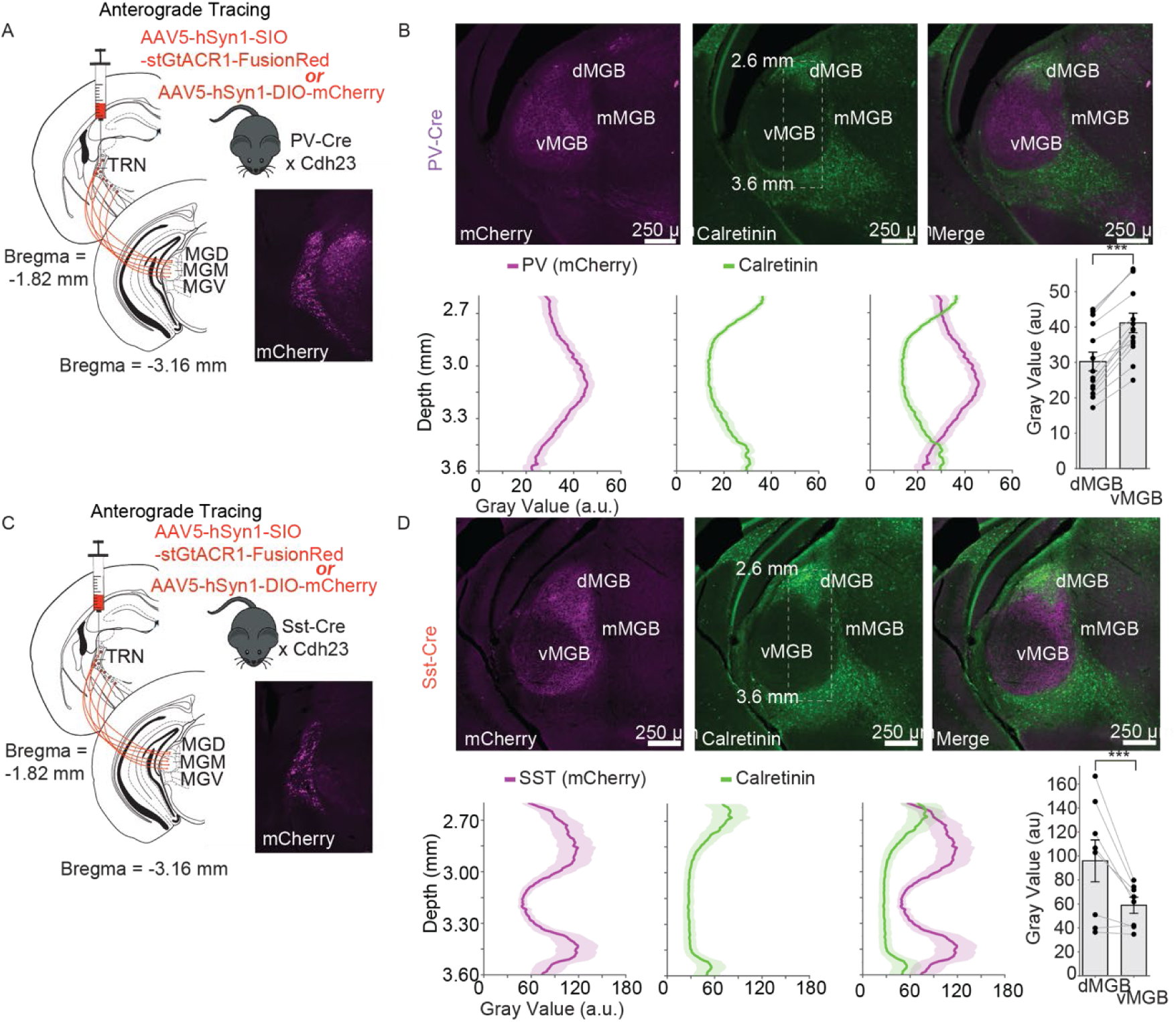
PV^TRN^ and SST^TRN^ project to different subregions of the MGB. **A.** Experimental design for anatomical experiments. PV-Cre mice were injected with AAV5-hSyn1-DIO-mCherry into TRN for anterograde tracing of PV^TRN^ neurons. Inset: mCherry expression in the TRN indicating the localization of the virus. **B. Top**: Projections of PV^TRN^ neurons primarily target the ventral subdivision of the MGB. *Left panel*: Viral expression of projections to the MGB (magenta). *Middle panel*: Immunohistochemical labeling of calretinin in the MGB (green). *Right panel*: Merged channels. **Bottom**: Quantification of the fluorescence intensity of projections from PV^TRN^ neurons that target MGB (Dashed box in top middle panel outlining the ROI). *Left panel*: intensity of viral expression of projections to the MGB in the ROI along the dorso-ventral axis. Dashed lines indicate extent of vMGB. *Middle panel*: intensity of the immunohistochemical labeling of calretinin in the MGB in the ROI along the dorso-ventral axis. *Right panel*: merged channels. *Bar plot:* mean intensity of the projections of PV^TRN^ neurons in dorsal and ventral MGB (N =14 ROIs; n = 3 mice). **C.** SST-Cre mice were injected with AAV5-hSyn1-DIO-mCherry into TRN for anterograde tracing of SST^TRN^ neurons. **D. Top**: Projections of SST^TRN^ neurons primarily avoid the ventral subdivision of the MGB. *Left panel*: Viral expression of projections to the MGB (magenta). *Middle panel*: Immunohistochemical labeling of calretinin in the MGB (green). *Right panel*: Merged channels. **Bottom**: Quantification of the fluorescence intensity of projections from SST^TRN^ neurons that target MGB (Dashed box in top middle panel outlining the ROI). *Left panel*: intensity of viral expression of projections to the MGB in the ROI along the dorso-ventral axis. *Middle panel*: intensity of the immunohistochemical labeling of calretinin in the MGB in the ROI along the dorso-ventral axis. *Right panel*: merged channels. *Bar plot:* mean intensity of the projections of PV^TRN^ neurons in dorsal and ventral MGB (N = 8 ROIs; n = 3 mice). *Scale bar = 0.25 mm. vMGB: ventral MGB; dMGB: dorsal MGB; mMGB: medial MGB; White dashed box in the top middle panel indicates ROI for fluorescence intensity quantification*.

### Inactivation of PV^TRN^ and SST^TRN^ neurons differentially affects tone-evoked responses in the MGB

Motivated by the anatomical distinctions described above and previous work identifying genetically distinct TRN neurons that express different anatomical and physiological properties^42,43^, we next tested whether and how sound responses of MGB neurons are modulated by PV^TRN^ and SST^TRN^ inputs. We injected a Cre-dependent soma-targeted inhibitory opsin (AAV5-hSyn-SIO-stGtACR1-FusionRed) into the audTRN of SST-Cre or PV-Cre mice (**Figure 2A-B, 2G-H, Supplementary Figure S1**). This virus only expresses in the soma of Cre-expressing neurons, reducing the chance of effects from backpropagating action potentials from local terminals^47^, and because TRN in mice is the only major locus of PV or SST expression close to the injection target, off-target impacts of light are minimized in this experiment. In separate experiments, we verified that the virus expression was restricted to the appropriate cell type (**Supplementary Figure S1**). We then implanted an optic fiber above audTRN, a headpost, and a ground pin for awake *in-vivo* electrophysiological recordings. In our experiments, mice passively listened to 50-ms tones ranging from 3 to 80 kHz (tone-On trials). In some tone-On trials we optogenetically inactivated PV^TRN^ or SST^TRN^ activity simultaneously with the tone (light-On trials), and we also presented light alone (light-Only trials) (**Figure 2A, G**). We recorded and measured the spiking activity of MGB neurons before, during and after tone and/or light presentations. After selecting for neurons that were sound-responsive in the 0-25 ms window after tone onset, we compared firing rates for the light-Off and light-On conditions of tone presentation and identified both significantly facilitated and suppressed MGB responses (**Figure 2B, C, H, I**). We then visualized the magnitudes of both types of modulated responses as a function of probe depth within the MGB (**Figure 2D-F, 2J-L**). To confirm that our optogenetic manipulations indeed inhibited audTRN neurons, in a separate set of experiments we tested and found that light stimuli over the TRN suppressed tone-evoked neuronal activity in neurons recorded in TRN (**Supplementary Figure S2**). We also verified that light stimuli suppressed spiking in whole-cell recordings of neurons expressing stGtACR1 in slices (**Supplementary Figure S3**).

**Figure 2.**
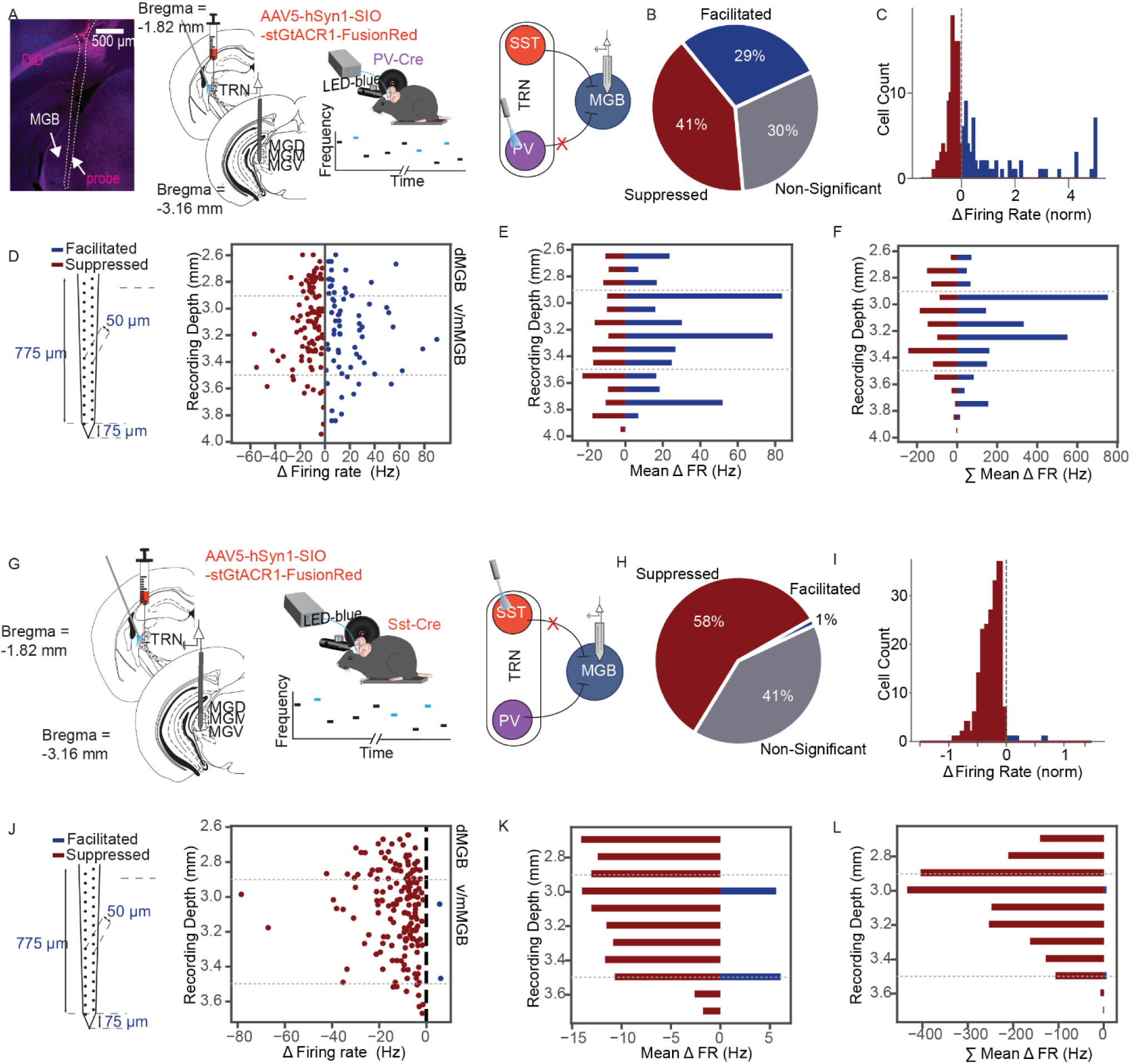
Inactivation of PV^TRN^ and SST^TRN^ neurons differentially affect tone-evoked activity in MGB neurons. **A.** PV-Cre mice were injected with AAV5-hSyn1-DIO-stGtACR1-FusionRed into the audTRN and were implanted with an optic fiber. We recorded from MGB neurons in awake head-fixed mice. We presented a set of pure tones in light-On and light-Off conditions. We selectively inactivated PV^TRN^ neurons while recording from neurons in the MGB using a vertical multi-channel electrode that spanned the depth of dMGB and vMGB, with the top electrode positioned at the tip of dMGB. Left: Example probe track labeled using DiD (magenta), from a recording in MGB. Center: Images from Paxinos Mouse Brain Atlas with target injection and recording locations superimposed. Right: Diagram of stimuli and recording setup. **B.** Pie chart breaking down the effect of PV^TRN^ inactivation on neurons of the MGB. We found that 41% of recorded units had suppressed tone-evoked responses (red slice), 30% of recorded units had facilitated tone-evoked responses (navy blue slice), and 29% of tone-responsive units were not affected by light manipulation (gray slice). **C.** Distribution of significant modulations of MGB responses during inactivation of PV^TRN^ neurons. Red: suppressed neurons; Blue: facilitated neurons. **D.** Scatter plot showing the change in firing rate (FR) of recorded MGB neurons with PV^TRN^ inactivation as a function of probe depth. Dotted grey lines indicate MGB subregion borders. **E.** Mean change in FR of recorded MGB neurons with PV^TRN^ inactivation as a function of probe depth in 100 mm bins. **F.** Total change in FR of recorded MGB neurons with PV^TRN^ inactivation as a function of probe depth in 100 mm bins. **G-L.** Same as **A-F**, but for SST^TRN^ inactivation, where we found that 58% of recorded units had suppressed tone-evoked responses (red slice), 1% of recorded units had facilitated tone-evoked responses (navy blue slice), and 41% of tone-responsive units were not affected by SST^TRN^ inactivation (gray slice). (N (PV) = 256, n = 4 mice. N (SST): 312, n = 4 mice). ROIs; n = 3 mice). *Scale bar = 0.50 mm. vMGB: ventral MGB; dMGB: dorsal MGB; mMGB: medial MGB*.

Because TRN is a powerful inhibitor of MGB, we initially expected to find that inactivating TRN neurons would drive disinhibition of sound-driven responses in MGB neurons. We expected this effect to be strongest in central vMGB, the primary sound relay, and mediated by the inhibitory PV^TRN^ neurons that innervate it. Alternatively, because neurons within TRN might be connected by reciprocal inhibition, inactivation of one subset of TRN neurons could disinhibit of another subset of TRN neurons and result in doubled inhibition, or simply, increased suppression of sound responses in MGB.

Indeed, inactivating PV^TRN^ neurons facilitated sound-evoked responses in a large fraction of MGB neurons (**Figure 2B, C**). The facilitation was stronger for the ventral regions of MGB than dorsal MGB (*p =* 0.008, binned by depth: 2.5-2.9 mm versus 3.1-3.5 mm, Mann-Whitney U test), and was exacerbated by the larger number of facilitated neurons in center (**Figure 2D-F**). Surprisingly, there was also a strong suppression of a fraction of neurons throughout MGB. However, this suppression was not differential across the depth of MGB (*p =* 0.219, Mann-Whitney U test). Inactivating SST^TRN^ neurons resulted in a dominant suppression of tone-evoked responses in MGB, with a handful of neurons exhibiting facilitation (**Figure 2H, I**). Interestingly, this suppression was also stronger in the dMGB than vMGB (*p =* 0.012, Mann-Whitney U test) (**Figure 2J-L**). The 4 cells that facilitated were in the v/mMGB (**Figure 2J**). Together with our anatomical results, these findings suggest that PV^TRN^ neurons provide direct inhibition or disinhibition to MGB, while SST^TRN^ neurons predominantly disinhibit the thalamocortical auditory pathway.

### Inactivation of PV^TRN^ neurons facilitates or suppresses tone-evoked responses in MGB

We next examined in more detail the effect of inactivating PV^TRN^ neurons on tone-evoked responses in MGB. For a representative *facilitated* MGB neuron, the raster plots and peri-stimulus time histogram (PSTH) (**Figure 3A-B**) demonstrate an increase in activity during tone presentation. For this neuron, the increase was most pronounced in the first 25 ms after stimulus onset and affected the responses across tone frequencies (**Figure 3C**). For a representative *suppressed* MGB neuron, the raster plot and PSTH depict a reduction in spiking activity during the tone (**Figure 3D-F**) that was most pronounced in the first 25 ms after tone onset.

**Figure 3.**
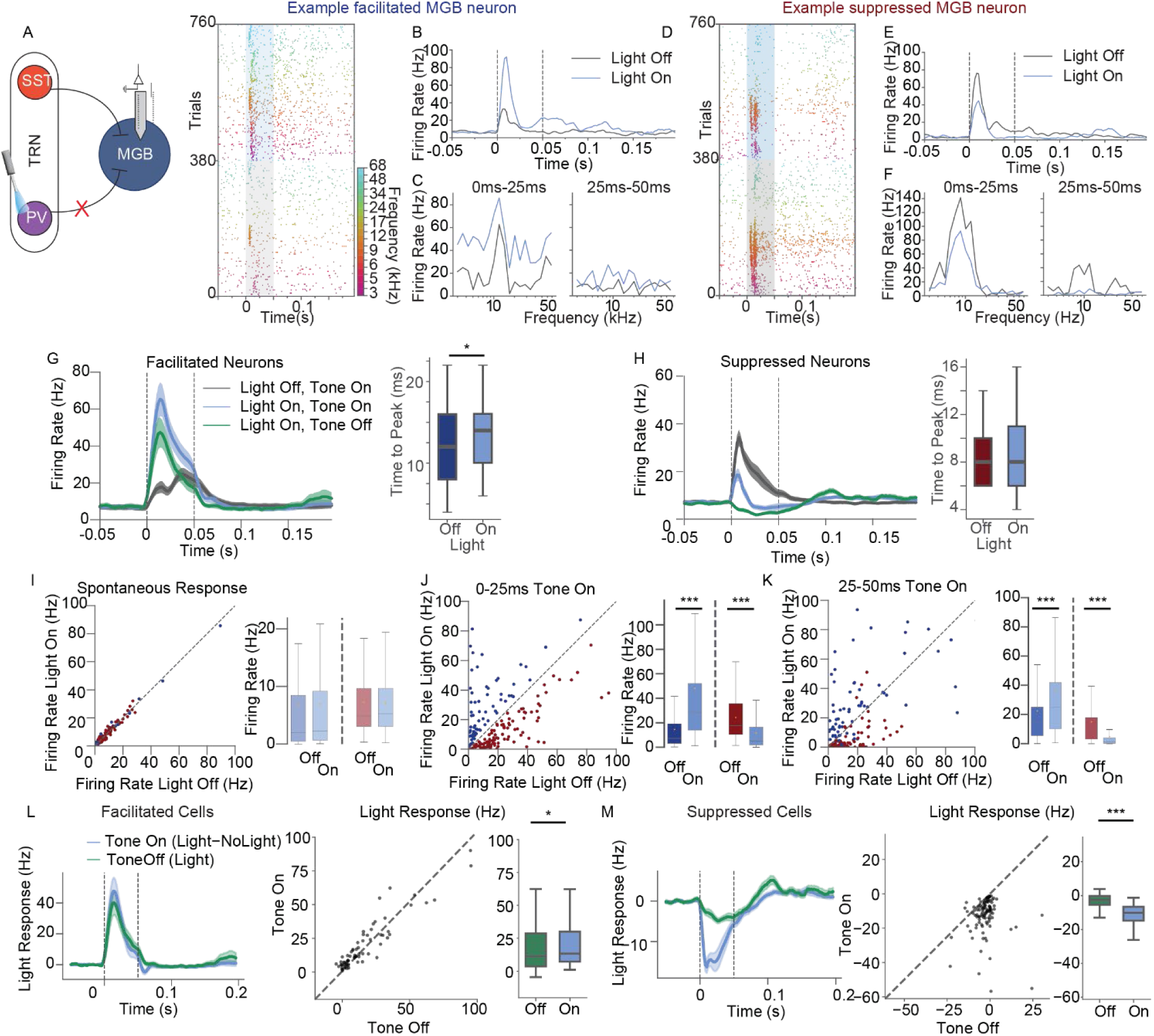
Inactivation of PV^TRN^ neurons facilitates or suppresses tone-evoked activity in MGB neurons. **A.** PV-Cre mice were injected with AAV5-hSyn1-DIO-stGtACR1-FusionRed into the audTRN and were implanted with an optic fiber. We selectively inactivated PV^TRN^ neurons on half of the trials while recording from MGB neurons in awake head-fixed mice in response to while a pure tones. In light-On trials, light was presented simultaneously with the tone. Raster plot of an example facilitated unit; trials are ordered by frequency and light condition. The gray box indicates light-Off conditions, and the blue box indicates the light-On conditions. **B.** PSTH of the facilitated example unit averaged across all trials for light-Off (black line) and light-On (blue line) conditions. **C.** Frequency response function for the facilitated unit during the early and late 25 ms of tone presentation during light-Off and light-On conditions. **D-F.** same as A-C, but for a representative suppressed neuron. **G, H.** Left: Mean PSTH of facilitated (G) and suppressed (H) recorded neurons in MGB (N = 74 facilitated and N = 104 suppressed neurons) for Light-Only trials (green line), tone-On, light-Off trials (black line), and tone-On, light-On trials (light blue line). Right: Times to peak of tone-evoked responses for facilitated (G, blue) and suppressed H, red) MGB neurons for light-Off (blue/red) and light-On (light blue) trials. **I-K.** Left: scatter plots and box plots of the firing rates for light-Off and light-On conditions for facilitated (blue) and suppressed (red) neurons during spontaneous activity prior to tone/light presentation (**I**), 0-25 ms post tone onset (**J**), and 25-50 ms post tone onset (**K**). **L, M.** Left panels: baseline-subtracted light responses for light-Only stimuli (green) , and light-induced change in firing rate between the two tone-On conditions (blue, difference between tone-On, light-On and tone-On, light-Off firing rates). Middle panel: scatter plot, and right panel: Box plots of mean firing rate changes in light response in tone-On vs. tone-Off conditions for the first 0-25 ms of stimulus presentation. **L:** facilitated neurons. **M:** suppressed neurons. Dashed line is x = y. *N = 256 units; n = 4 mice. Here and below, boxplots depict the median and interquartile range (IQR) for each condition; whiskers extend to values within 1.5xIQR; shaded areas in psths represent SEM±; error bars represent SEM±. *** denotes p-value < 0.001; * denotes p-value < 0.05*.

Across the population of MGB neurons, PV^TRN^ inactivation during tone presentation resulted in facilitation in 30% of neurons; 41% of neurons were suppressed ,and 29% of neurons in the MGB were not significantly affected by PV^TRN^ inactivation (**Figure 2B, C**). Responses of facilitated neurons demonstrated a sharp peak followed by a prolonged decay in firing rate after tone onset in both light-Off and light-On conditions, and the peak of the tone responses was greater in the presence of light (**Figure 3G**). Response peaks of suppressed neurons to tones followed a similar timecourse but were reduced during light-On relative to tone-On, light-Off trials (**Figure 3H**). When PV^TRN^ neurons were inactivated, time-to-peak significantly increased in facilitated MGB neurons but was unchanged in suppressed MGB neurons (**Figure 3G, H right;** *p*(facilitated) < 0.05, *p*(suppressed) = 0.070; signed-rank test), suggesting that PV^TRN^ neurons selectively control dynamics of neuronal responses in MGB.

We verified that over the population, there were no significant differences in spontaneous activity prior to stimuli between the units that were later suppressed or facilitated by PV^TRN^ inactivation during tone presentation (**Figure 3I**; *p* = 0.856, t-test). Facilitation or suppression of mean tone-evoked firing during the first 25 ms of the tone by inactivation of PV^TRN^ was significant over the population (**Figure 3J**, 0-25 ms post tone onset) and those effects persisted for the duration of the tone (**Figure 3K**, 25-50 ms post tone onset; N(facilitated): 74, N(suppressed): 104; *p*(facilitated) < 0.001, *p*(suppressed) < 0.001 for both time periods tested; signed-rank test).

To examine whether light responses in MGB – that is, the impacts of inactivating PV^TRN^ were affected by the tone, we subtracted the tone responses in the light-On and light-Off conditions. We then compared that difference to baseline-subtracted responses during light-Only (tone-Off) trials. In facilitated neurons, light responses were larger in the tone-On condition than responses to the light-Only (tone-Off) condition (**Figure 3L**, N = 74, *p* = 0.011). In suppressed neurons, light caused a greater suppression of firing in the tone-ON condition compared to the light-Only condition (**Figure 3M**, *N =* 104, *p* < 0.0001). These differences in light responses suggest that inhibition provided by PV^TRN^ neurons is supralinear – it is stronger in the presence of tone than in silence and may reflect sound responses of TRN neurons.

We also asked tested whether the effects of PV^TRN^ inactivation differed for MGB recording site(s) that were tonotopically organized along the probe depth (dorsal-ventral axis, increasing best frequency with depth) and those that were not. We found only small differences between the effects of PV^TRN^ inactivation with a slightly greater fraction of neurons suppressed in tonotopic than non-tonotopic sites (**Supplementary Figure S4**).

Overall, these results demonstrate that inactivating PV^TRN^ neurons rapidly and bi-drectionally affected tone-evoked responses in MGB. Thus, PV^TRN^ neurons can both directly suppress neuronal responses in MGB or enhance the response firing rate, thereby involving multiple mechanisms that shape MGB firing rates.

### Inactivation of SST^TRN^ neurons primarily suppresses tone-evoked responses of MGB neurons

We next examined in more detail the effect of inactivating SST^TRN^ neurons on tone-evoked responses of MGB neurons. For a representative *facilitated* MGB neuron, the raster plots and post-stimulus time histograms (**Figure 4A-B**) demonstrate an increase in activity at the onset of tone presentation. For this neuron, the increase was present in the first 25 ms after stimulus onset and affected the responses to frequencies near the neuron’s best frequency (**Figure 4C**). For a representative *suppressed* MGB neuron, the raster plot and PSTH depict a reduction in spiking activity throughout the tone (**Figure 4D, E**). The suppressive effects were most pronounced in the first 25 ms after tone onset (**Figure 4F**).

**Figure 4.**
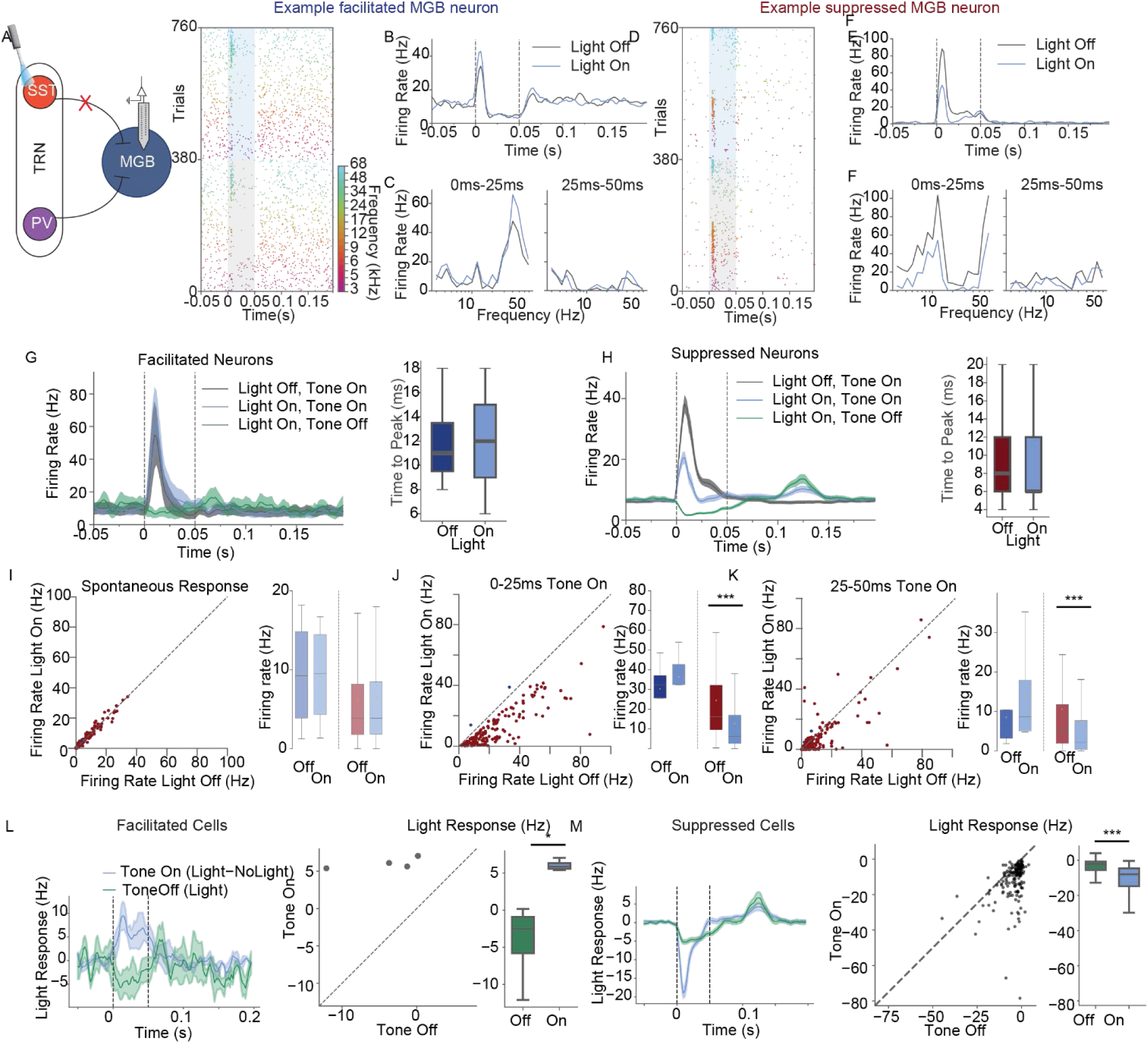
Inactivation of SST^TRN^ neurons predominantly suppresses tone-evoked activity in MGB neurons. **A.** SST-Cre mice were injected with AAV5-hSyn1-DIO-stGtACR1-FusionRed into the audTRN and were implanted with an optic fiber. We recorded from MGB neurons in awake head-fixed mice. We presented a set of pure tones in light-On and light-Off conditions. Light was simultaneous with the tone. We selectively inactivated SST^TRN^ neurons while recording from neurons in MGB. Raster plot of an example facilitated unit; trials are ordered by frequency and light condition. The gray box indicates light-Off conditions, and the blue box indicates light-On conditions. **B.** PSTH of the facilitated example unit averaged across all trials for the light-Off (black line) and light-On (blue line) conditions. **C.** Frequency response function for the facilitated unit during the early and late 25 ms of tone presentation during light-Off and light-On conditions. **D-F.** same as A-C, but for a representative suppressed neuron. **G, H.** Left: Mean PSTH of facilitated (G) and suppressed (H) recorded neurons in MGB (N = 4 facilitated. N = 181 suppressed) for light-Only trials (green line), tone-On, light-Off trials (black line), and tone-On, light-On trials (light blue line). Right: Time to peak of tone-evoked responses for facilitated (G, blue) and suppressed (H, red) MGB neurons for light-Off (blue/red) and light-On (light blue) trials. **I-K.** Scatter plots and box plots of mean firing rates for light-Off and light-On conditions for facilitated (blue) and suppressed (red) neurons during spontaneous activity (**I**), 0-25 ms post stimulus onset (**J**), and 25-50 ms post stimulus onset (**K**). **L, M.** Left panels: baseline-subtracted light responses for light-Only stimuli (green), and light-induced change in firing rate between the two tone-On conditions (blue, difference between tone-On, light-On and tone-On, light-Off firing rates). Center: scatter plot of firing rate changes in light response in tone-On vs. tone-Off conditions. Right: box plot of mean light responses in tone-Off and tone-On conditions. Dashed line is x = y. **L:** Facilitated neurons. **M:** Suppressed neurons. *N = 312 units; n = 4 mice*.

Across the population of neurons, during tone presentation and SST^TRN^ inactivation 58% neurons were suppressed, only 1% of neurons were facilitated and 41% of neurons in the MGB were not affected by SST^TRN^ inactivation (**Figure 2H, I**). The average PSTH across the few facilitated neurons (N = 4 neurons) shows a small increase in firing during light presentation (**Figure 4G**). Suppression driven by SST^TRN^ neuronal inactivation followed a similar timecourse as the effects of PV^TRN^ inactivation (**Figure 4H**). There was no change in the time-to-peak response of the 4 facilitated MGB neurons, and no change for time to peak in suppressed MGB neurons, upon inactivation of SST^TRN^ neurons (**Figure 4G, H right;** *p*(facilitated) = 1.0, *p*(suppressed) = 0.071; signed-rank test).

We verified that over the population, there were no significant differences in spontaneous activity prior to stimuli across units that were later suppressed or facilitated by SST^TRN^ inactivation during tone presentation (**Figure 4I**; *p* = 0.458, t-test). In neurons classified as suppressed, neuronal activity was significantly reduced by SST^TRN^ inactivation during the tone presentation for both the first 25 ms (**Figure 4J**) and the second 25 ms (**Figure 4K**) of tone presentations, while changes in the small set of facilitated neurons were not significant as a group (**Figure J, K;** N(suppressed): 181, N(facilitated): 4; *p*(facilitated) =0.125, *p*(suppressed) < 0.001 for both time windows; signed-rank test). We found that SST^TRN^ inactivation suppressed a greater fraction of neurons in non-tonotopically than in tonotopically organized recording sites, suggesting that the disinhibition provided by SST^TRN^ neurons may be localized to non-lemniscal MGB subdivisions consistent with our anatomical findings (**Supplementary Figure S5**).

To examine whether light responses in MGB – that is, the impacts of inactivating SST^TRN^ were affected by the tone, we compared the baseline-subtracted light-Only response to the difference in tone-On responses during light-On and light-Off trials. The response to the light was increased during the tone in the few facilitated neurons in comparison to the light-only response (**Figure 4L**, *N =* 4, *p* = 0.026), and decreased in suppressed neurons (**Figure 4M**, *N* = 181, *p* < 0.0001), again suggesting a supralinear effect of SST inhibition during tone presentation and possible contributions from sound responses of TRN neurons. Overall, these results demonstrate that inactivating SST^TRN^ neurons largely drives suppression of MGB neurons.

### PV^TRN^ and SST^TRN^ differentially affect frequency tuning of MGB neurons

Inactivation of PV^TRN^ neurons produced significant changes in MGB frequency tuning for both the facilitated- and suppressed- <GB neuronal populations, with a significant shift upwards and a significant shift downwards in the frequency response functions, respectively (**Figure 5A-B**; N(facilitated): 74, N(suppressed): 104; *p*(facilitated-all octaves) < 0.001, *p*(suppressed-all octaves) < 0.001; signed-rank test). To examine frequency selectivity of neurons (or how specific neuronal responses are to a best frequency) we calculated the sparseness index across the responses. In the sparseness index, a value close to 1 indicates that units respond more selectively to a preferred stimulus or central frequency, whereas a value close to 0 indicates that units respond to a broader range of stimuli or frequencies. We found that inactivation of PV^TRN^ significantly affected sparseness in both facilitated and suppressed subgroups. Following PV^TRN^ inactivation, sparseness in facilitated MGB neurons significantly decreased, indicating that neurons became more broadly responsive (**Figure 5A**; *p* < 0.001; signed-rank test). Interestingly, in suppressed MGB neurons, PV^TRN^ inactivation increased sparseness, indicating that neurons on average became more narrowly responsive (**Figure 5B**; p < 0.001; signed-rank test). These results indicate that PV^TRN^ neurons contribute to how narrowly or broadly tuned MGB neurons are to auditory stimuli.

**Figure 5.**
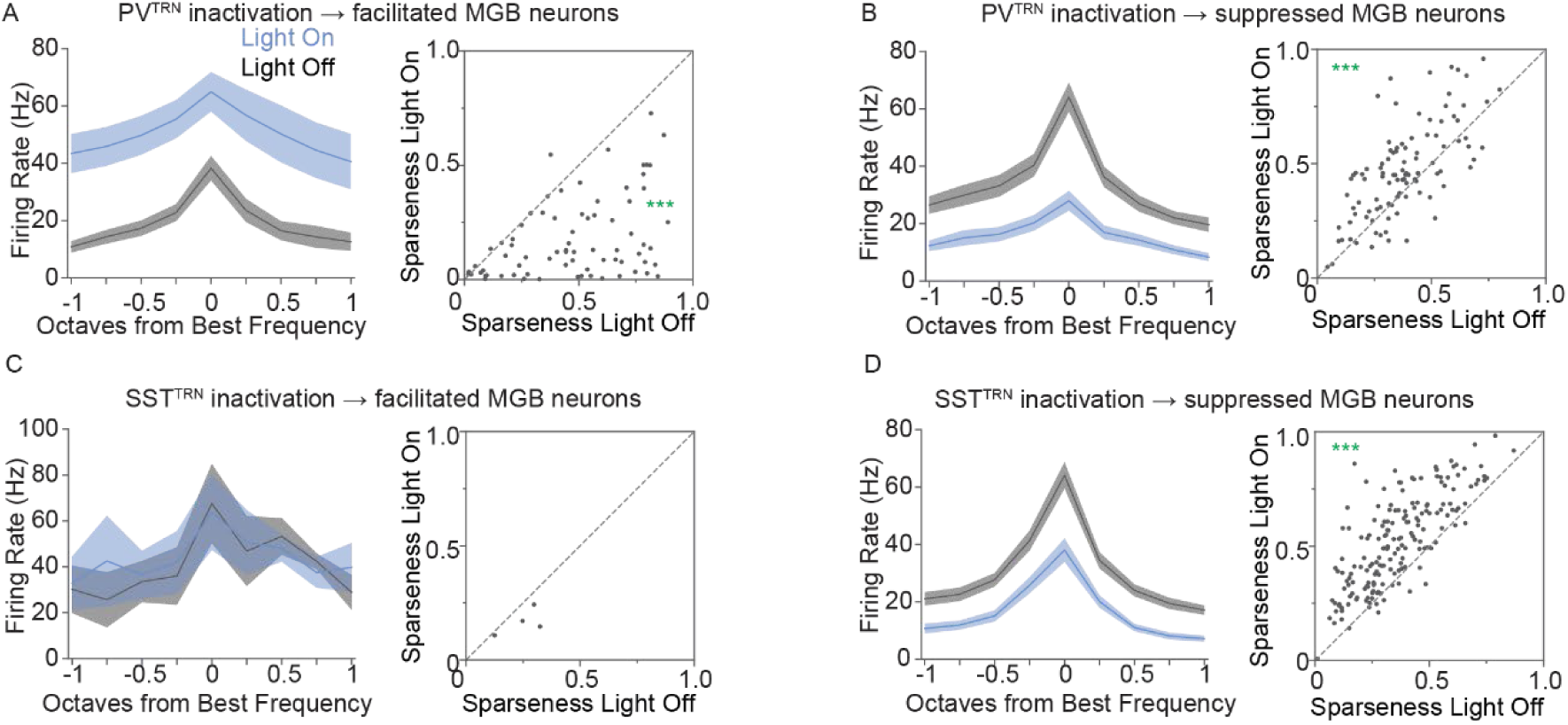
Inactivation of PV^TRN^ and SST^TRN^ neurons differentially affect frequency tuning of MGB neurons. A-D. *Left:* Mean frequency response functions for light-On (light blue) and light-Off (light gray) trials for neurons, centered on best frequency (BF = 0 octaves). Shaded areas represent SEM±. *Right.* Scatter plots of sparseness for light-On versus light-Off conditions. **A, C.** Facilitated neurons. **B, D.** Suppressed neurons. *N = 256 neurons; n = 4 mice (PV). N = 312 neurons; n =4 mice (SST). *** denotes p-value < 0.001; * denotes p-value < 0.05*.

Inhibition of SST^TRN^ neurons produced no significant changes in tuning for the few MGB neurons that had facilitated tone-evoked responses (**Figure 5C**; *N* (facilitated): 4; *p* > 0.1; signed-rank test). However, in the MGB neurons suppressed by SST^TRN^ inactivation, firing rates showed a significant downward shift in the frequency response function (**Figure 5D**; *N* = (suppressed): 181; *p* < 0.001; signed-rank test). Inactivation of SST^TRN^ neurons did not affect sparseness in the small facilitated group (**Figure 5C***; N* (facilitated): 4; *p* = 0.125; signed-rank test). However, similar to PV^TRN^ inactivation, sparseness in suppressed MGB neurons increased significantly during SST^TRN^ inactivation (**Figure 5D**; *N* (suppressed): 181; *p* < 0.001; signed-rank test).

We also examined whether the best frequency and tuning width for frequency-tuned MGB neurons changed during PV^TRN^ or SST^TRN^ inactivation. To do this, we fit a Gaussian function to each MGB neuron tuning curve in both the light-On and light-Off conditions (examples, **Supplementary Figure 6 D, H, L**) and compared the standard deviation σ parameter, which corresponds to the width of frequency tuning. In line with our findings that PV^TRN^ and SST^TRN^ inactivation increased sparseness in suppressed MGB neurons, MGB neurons that were suppressed by PV^TRN^ inactivation showed a significant decrease in σ (PV: **Supplementary Figure S6 F, G**, N = 26, t = 2.42, p = 0.027; SST: **Supplementary Figure S6 J, K,** N = 47, t = 3.30, p = 0.0002). However, for neurons facilitated by PV^TRN^ inactivation, there was no change in σ between light-On and light-Off conditions (**Supplementary Figure S6 A, B,** N = 11, t = -1.27, p = 0.278) and there were no well-fit facilitated neurons during SST^TRN^ inactivation. Together, these results suggest that changes in MGB response sparseness may at least partially be due to changes in the width of frequency response function evoked by manipulation of TRN inhibitory neurons.

### Specifics of connectivity between MGB-TRN circuits determine the sign of MGB responses during TRN cell-type inactivation

To examine the many possible connectivity schemes of primary and higher-order auditory thalamocortical circuits that may account for the observed changes in sensory responses during inactivation of specific TRN populations, we constructed a 4-cell Hodgkin-Huxley model (see Methods) comprising two channels: a primary vMGB relay cell reciprocally connected to a PV^TRN^ cell, and a higher-order MGB cell reciprocally connected to a SST^TRN^ cell (**Figure 6A-B**), matching the subtype-specific connections that we identified and have been previously demonstrated for other sensory modalities^42,43^. Without connectivity between primary and higher-order channels, inactivation of the PV^TRN^ cell always allowed substantially increased responses in the vMGB firing rate to a tone input, whereas inhibiting the SST^TRN^ cell has no effect at all on vMGB response, as expected (not shown). Those trials served as a comparison control for subsequent trials.

**Figure 6.**
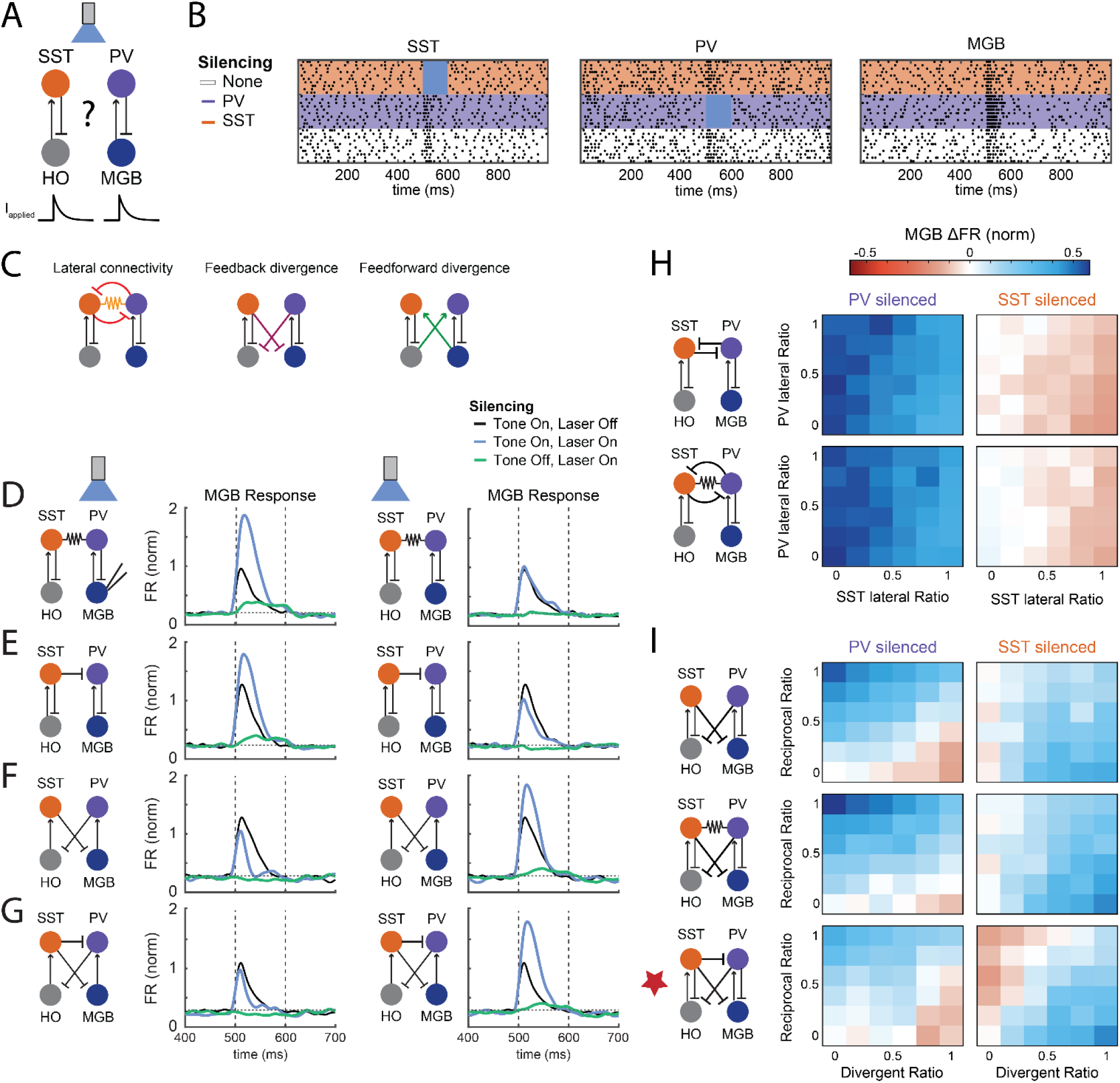
Schemes of circuit connectivity that account for bidirectional modulation of vMGB responses during subtype-specific inactivation in TRN. **A**. 4-cell model of primary sensory MGB-TRN pairs and higher order (HO) MGB-TRN pairs. **B.** Raster plots from trials with no inactivation, during PV inactivation (purple) and during SST^TRN^ inactivation (orange). **C.** Candidate connectivity schemes tested in models. **D.** PSTH of MGB firing for circuits with electrical coupling between TRN subtypes. MGB responses are normalized to the peak response during non-inactivated trials with no connectivity between columns. Left, primary MGB firing rate to input alone (black line) and with PV^TRN^ silencing (light blue line). Right, MGB firing rate input alone (black line) and with SST^TRN^ silencing (light blue line). **E**. As in D, for a circuit including lateral SST^TRN^ to PV^TRN^ connections. **F**. Effect of inactivation on MGB rate for heterogeneous combinations of lateral inhibitory connections, with and without electrical synapses between TRN cells. The ratio of lateral synapses is the percent of trials that included lateral synapses for each TRN cell type. **G.** As in D, for circuits with both reciprocal and divergent feedback inhibition. **H.** As in D, for circuits containing only divergent inhibition. **I.** Effect of cell type-specific inactivation on MGB rates for heterogeneous inhibitory feedback connections between TRN and thalamic cells. Divergence ratio is the percent of trials that included divergent synapses. Reciprocal ratio is the percent of trials that included reciprocal TRN-MGB synapses. Middle, with electrical coupling between PV and SST cells. Bottom, effects of TRN inactivation on MGB rate with heterogeneous inhibitory connections in addition to lateral inhibition from the SST^TRN^ to PV^TRN^ cells.

We first examined whether intra-TRN connectivity would suffice to produce both suppression and facilitation of vMGB responses during cell type-specific silencing (**Figures 3,4**). Electrical synapses extensively couple TRN cells, including coupling between subtypes^45^, that may substantially modulate activity within the thalamocortical circuits examined here^48,49^. We examined the effect of inter-type TRN coupling, using a strong electrical synapse with coupling coefficient ∼0.2, on the responses of the primary MGB cell. Similar to the uncoupled control case, vMGB responses were facilitated when PV^TRN^ was inactivated (**Figure 6D**, left), and once again as expected we saw no changes in vMGB responses during inactivation of SST^TRN^ (**Figure 6D, right**). Although the existence of inhibitory synapses between TRN neurons in mature tissue has been controversial ^50^, there is some evidence for lateral inhibition within TRN^51,52^. Inclusion of a lateral inhibitory synapse from SST^TRN^ to PV^TRN^ allowed for facilitated vMGB response during PV^TRN^ inactivation, and we noted that MGB responses during no-inactivation trials were also elevated from the control response (**Figure 6E, left**). Silencing SST^TRN^ cells in this circuit produced a substantial reduction in the tone response of the vMGB neuron compared to control responses (**Figure 6E, right**), matching the experimental results (**Figure 2, 4**) through relief of disynaptic inhibition. We also noted changes in the spontaneous rates for light-Only trials during subtype silencing that was consistent with the experimental recordings (**Figure 6E, cyan lines**). Silencing either of the cell types in a circuit that included inhibitory connections from PV^TRN^ to SST^TRN^ did not reduce tone responses of vMGB, however (not shown). Thus, one specific form of intra-TRN connectivity, from SST^TRN^ to PV^TRN^ cells, is a candidate circuit mechanism underlying the unexpected suppression of MGB during cell type-specific TRN inactivation.

Next, we asked whether the responses of primary vMGB change during cell type-specific inactivation in the context of variations in feedback inhibition. TRN cells send divergent inhibitory connections to several MGB cells but do not often send inhibition back to the same cells that excite them^53,54^. Additionally, activation of one thalamic nucleus can produce inhibition in nearby sensory-related nuclei (i.e. activation of POM evoking IPSCs at VPM) thought to arise from the TRN^55,56^. Thus, we included and varied the amount of divergent feedback inhibition from the TRN to the MGB in our circuits. In circuits with only divergent but lacking reciprocal inhibitory feedback, PV^TRN^ inactivation produced a suppression of vMGB responses (**Figure 6F, left**), while SST^TRN^ inactivation produced facilitation of vMGB responses (**Figure 6F, right**). In circuits with only divergent inhibitory feedback as well as a lateral SST^TRN^ to PV^TRN^ connection, PV^TRN^ inactivation produced a suppression of vMGB responses (**Figure 6G, left**), while SST^TRN^ inactivation produced greater facilitation of vMGB responses (**Figure 6G, right**). Thus, we conclude that uniformly divergent feedback inhibition is unlikely to account for the experimental observations of vMGB suppression during SST^TRN^ inactivation.

As we anticipate that spectrums of inhibitory connectivity and strength may co-exist in live tissue, we then asked whether the relative strengths of within-TRN, lateral inhibition and TRN-MGB reciprocal inhibition could control the sign of vMGB responses during selective silencing. To test the effect of heterogeneous circuitry on firing rate changes in primary vMGB cells, we varied the ratio of lateralization within the modeled circuits over sets of trials. In circuits where TRN and MGB were reciprocally connected, varied relative strengths of lateral intra-TRN inhibition always produced facilitation of vMGB responses when PV^TRN^ was inactivated (**Figure 6H, left**); the effect of SST^TRN^ lateralization was only to modulate the total facilitation produced. During SST^TRN^ inactivation, increases in SST^TRN^ lateral inhibition suppressed MGB responses independently of the presence of PV^TRN^ lateral inhibitory synapses (**Figure 6H, top right**). We note that the addition of coupling between TRN cells allowed for bidirectionality in changes in vMGB rate when SST^TRN^ was inactivated for networks with weak SST-to-PV lateralization (**Figure 6H, bottom right**). From this set of simulations addressing intra-TRN connectivity, we conclude that inactivating SST^TRN^ can reproduce the unexpected suppression of vMGB responses when SST-to-PV synapses or gap junctions are included in the circuit; however, these lateral intra-TRN connections alone cannot reproduce the bidirectional effects of PV^TRN^ inactivation.

Finally, we asked if variations in the relative power of the two types of feedback inhibition, combined, could allow both vMGB enhancement and suppression during cell type-specific inactivation trials. Increasing the relative strength of the divergent inhibitory synapses decreased the responses of the vMGB cell, while increasing the strength of reciprocal inhibition increased those responses, during PV^TRN^ inactivation trials compared to the baseline firing rate (**Figure 6I, top left**). This is consistent with the bidirectionality of the experimental results for PV^TRN^ inactivation (**Figure 3**). SST^TRN^ inactivation produced only increases in vMGB activity, as divergent inhibitory ratio increased from zero (**Figure 6I, top left**), inconsistent with experiments. However, electrical coupling added to this circuitry allowed SST^TRN^ inactivation to diminish vMGB responses for weakly lateralized networks (**Figure 6I, middle**). Including SST^TRN^-to-PV^TRN^ lateral inhibition in this case modulated the SST^TRN^ inactivation response towards more suppressed outcomes (**Figure 6I, bottom right).** Together, our simulations lead us to conclude that divergent feedback connections, together with electrical synapses or SST^TRN^-to-PV^TRN^ inhibition, may be embedded within the thalamus and serve as the sources of the bidirectional vMGB responses observed during PV^TRN^ inactivation and the suppression during SST^TRN^ inactivation.

## DISCUSSION

Differences between intrinsic physiology, synaptic connections, and axonal projections of distinct cell types have been described for the GABAergic neurons of the TRN^42–45^, but the impact of those differences on their inhibitory modulation of thalamic processing remains unknown. We used a combination of viral anatomical tracing, optogenetics, awake *in-vivo* electrophysiological recording, and computational modeling to study the functional roles of audTRN PV and SST neurons in modulating sound responses in the MGB. Our results confirm that audTRN PV and SST neurons project to hierarchically distinct sensory thalamic regions: while PV^TRN^ neurons primarily project to the center of the primary or first-order auditory thalamic sub-region vMGB, SST^TRN^ neurons project almost exclusively to the dorsal subregions of MGB and higher-order auditory thalamus (**Figure 1**). Here, we performed recordings in awake mice to examine the effects of audTRN PV and SST neuron inactivation on the intact auditory circuit. We found that inactivating these distinct subpopulations of audTRN inhibitory neurons had diverging effects on auditory tone responses of MGB neurons **(Figure 2)**. While inactivation of PV^TRN^ neurons facilitated a large fraction of MGB neurons, we also found that inactivation of both PV^TRN^ and SST^TRN^ neurons suppressed a large fraction of responses in MGB (**Figures 2, 3, 4**). Our results show that inactivation of both PV^TRN^ and SST^TRN^ neurons altered the frequency tuning responses of MGB neurons **(Figure 5, S6)**. In addition, computational models allowed us to identify several embedded variations of circuitry between TRN and MGB that could enable the experimental observations (**Figure 6**, for summary see **Figure 7**).

**Figure 7.**
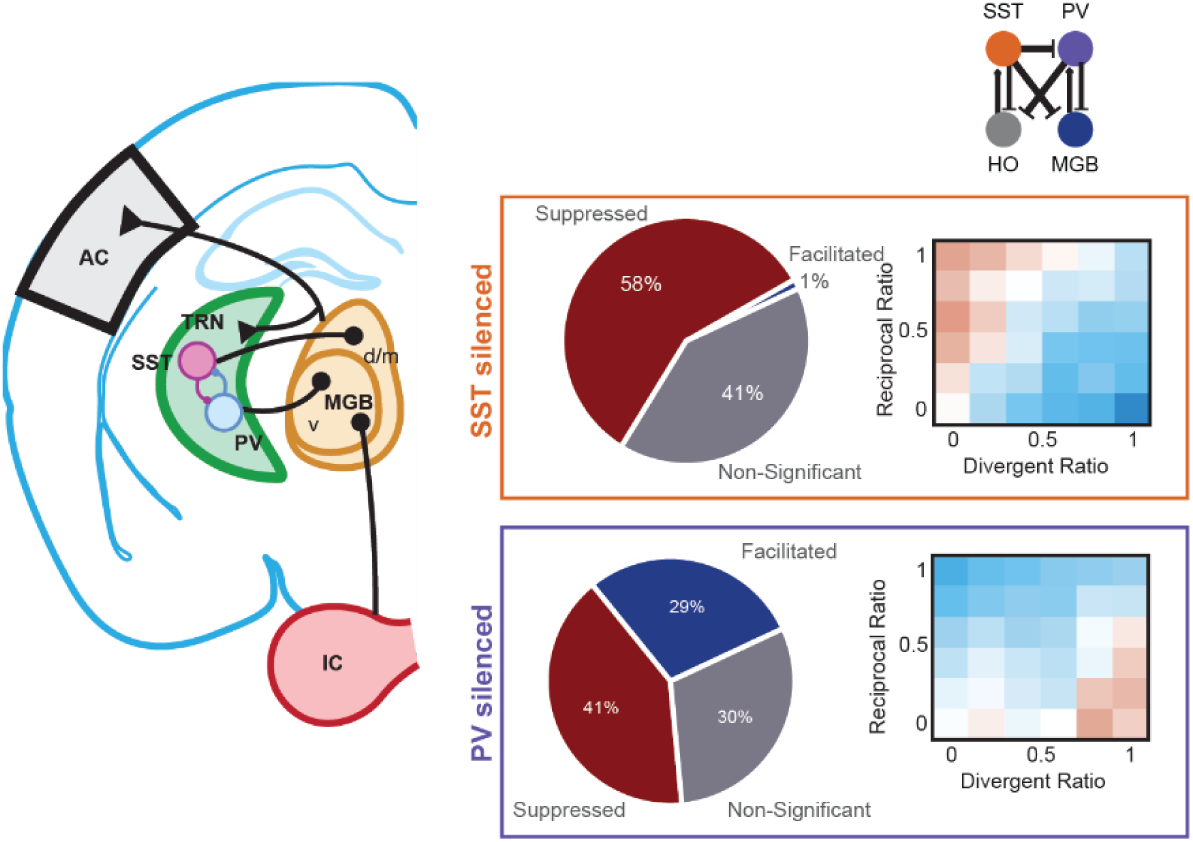
Inhibitory neurons differentially control sound-evoked responses in the MGB. Left: diagram of the proposed circuit underlying our results. Boxes: summary of effects of optogenetic manipulation of SST^TRN^ or PV^TRN^ inactivation on MGB firing and the circuitry and results of computational models that support a similar distribution of effects.

Together, our anatomical, electrophysiological, and simulation results indicate complex roles for TRN inhibition on MGB neurons and suggest a larger degree of divergence than what has been suggested from the anatomical work here and by others^42,43^. A variety of connection schemes have been proposed for TRN, including intra-TRN inhibition^51,51^, lack of intra-TRN inhibition^50^, typed reciprocal inhibition^42,43^, and open-loop or divergent connectivity to thalamic nuclei^55,56^. Cross-modal TRN responses have also been reported^57,58^, offering further evidence of nuanced connectivity. In addition, gap junctions of the TRN could be exclusive within one set or both sets of cell types or modality, or promiscuous across types^45^, or absent in adult tissue. Our computational survey of these possibilities suggests that divergence of TRN-MGB connectivity is necessary, in addition to reciprocal inhibition, to achieve the bidirectional responses in MGB observed for PV inactivation; and that some intra-TRN connectivity, whether gap junctional and/or GABAergic, plays a role in allowing for the suppressive effects of SST inactivation. Our modeling offers several predictions for future experiments using optogenetic approaches. First, our model predicts a cell-specific inhibitory connection from SST^TRN^ to PV^TRN^. Second, divergent inhibition (*e.g.* from SST^TRN^ to vMGB neurons) could also be evaluated in the same experiment. Moreover, our modeling predicts that SST^TRN^ are driven by higher order cells; thus inactivating the higher order population while also inactivating the SST^TRN^ would potentially reverse the observed impact on vMGB responses during SST silencing. Similar experiments in future studies could be applied to test a PV^TRN^-d/mMGB connection.

One proposed mechanism for our results is the possibility of intra-TRN synapses between PV^TRN^ and SST^TRN^ neurons, similar to the interactions in cortical regions^1^. Evidence for GABAergic synapses between nearby TRN neurons in slice preparations is exceedingly sparse^50,59^, but longer-range intra-TRN interactions are possible, and reports of dendro-dendritic synapses have been reported in the TRN of the cat^60^. Driven by our modeling results, our subsequent experiments found one example of such a connection in slice recordings (**Supplementary Figure S7**). It has also been reported that excitatory collateral inputs from thalamic relay nuclei drive excitatory input onto TRN neurons and therefore activate TRN neurons^61^. It is also possible that the suppression effects in tone-responsive units of the MGB is a direct result of fast feedback arising from excitatory MGB collaterals, intra-TRN dendro-dendritic synapses, or other regions of GABAergic inputs to the TRN like the basal forebrain, substantia nigra or lateral hypothalamus^62–65^.

While our experiments did not include a behavioral component, our results have clear implications for modulation from the TRN during active behaviors. Our results suggest that PV^TRN^ and SST^TRN^ neurons may play distinct roles in behaviors that rely on lemniscal or non-lemniscal thalamic function. The TRN receives diverse modulatory inputs, including projections from the substantia nigra reticulata^64^, the basal forebrain^65–67^, globus pallidus^68–70^ and hypothalamus^63^. Whether these inputs are cell type-specific is unknown. Our results indicate that the impact of inhibiting or modulating specific TRN neuron types is complex and raises interest in the types, sources, and targets of modulatory input to the thalamocortical pathways, along with the function of these projections. For example, inhibition of specific classes of TRN neurons that results in specific suppression or facilitation of the thalamocortical relay could be a mechanism for modulating sensory perception in line with behavioral state. Indeed, recent studies have shown the contributions of MGB to associative behavior memory tasks^71^.

Our experiments and analyses do not account for the feedback provided to MGB and TRN from the cortex. Within the auditory thalamus, the TRN provides the majority of inhibition to the MGB, however, MGB neurons also receive feedback from corticothalamic neurons in L5/L6 of AC as well as input from other association areas, such as the BLA^33,72,73^. It is possible that the suppressive effects we observed included a component of fast feedback inhibition from another region of the brain, or polysynaptic interactions. Future studies using high-density channel probes to record from multiple brain areas simultaneously could further our understanding of inhibitory circuit mechanisms. For example, recording from the TRN-MGB-AC circuit simultaneously while optogenetically manipulating 1) corticothalamic feedback, 2) TRN-MGB inhibitory synapses, or 3) collateral inputs from MGB to the TRN would allow for recording effects of audTRN on neural responses along the auditory pathway. An extensive anatomical tracing study would also specifically identify possible sources of GABAergic input to the PV^TRN^ and SST^TRN^ neurons involved in the auditory pathway. This could reveal possible sources of feedback inhibitory inputs to the TRN that might drive the tone-evoked suppression we observe in our study. In addition, descending cortical projections to the TRN are tonotopic, and this organization may result in PV^TRN^ and SST^TRN^ inhibition of the thalamocortical relay that is not only anatomically structured but also frequency specific. Future studies could examine how optogenetic inhibition of different frequency regions of auditory cortex alters the firing of PV^TRN^ and SST^TRN^ neurons, and the resulting changes in MGB neuron tuning curves.

Additionally, the role of the TRN in integrating sound information across the two ears remains to be investigated. In our study, viral injections and electrophysiological recordings were performed contralateral to the position of the speaker. Neurons in the MGB are biased toward the contralateral ear, and sound intensity from the freestanding speaker is stronger at the contralateral ear as opposed to the ipsilateral ear; therefore, we expected that the sound emitted by the speaker would predominantly drive neurons in the recorded location^74^. Nonetheless, sound still reaches the ear that is contralateral to the speaker, and that information is transferred to the contralateral superior olive and eventually the contralateral MGB (relative to the speaker) ^75^. Additionally, neurons in the MGB receive inputs from both ipsi- and contralateral IC^76^. In parallel, information about the sound is processed within the ipsilateral MGB relative to the speaker and inhibition from that ipsilateral TRN may affect processing in the contralateral MGB via direct or indirect projections. To disentangle these effects, future experiments may include reversible ear-plugging of the contralateral ear, bilateral neuronal modulations or simply varying the speaker position.

Combined, our results provide an insight into the complexity of the audTRN and identify cell-type specific mechanisms for controlling sound processing in the auditory thalamus.

## MATERIALS AND METHODS

### Animals

We performed all experimental procedures in accordance with NIH guidelines on use and care of laboratory animals and by approval of the Institutional Animal Care and Use Committee at the University of Pennsylvania (protocol number 803266). Care was taken to minimize any pain, discomfort, or distress of the animals during and following experiments. Experiments were performed in adult male (n = 12) and female (n = 12) mice aged 3-8 months and weighing 20-32g. The mouse lines used in this study were crosses between Cdh23 mice (B6.CAST-Cdh23^Ahl+^/Kjn; RRID:IMSR_JAX:002756)—a transgenic line that has a targeted point reversion in the Cdh23 gene which protects mice from age-related hearing loss^77^—and PV-Cre mice (B6.129P2-Pvalb^tm1(cre)Arbr^/J; RRID:IMSR_JAX:017320) or SST-Cre mice (SST^tm2.1(cre)Zjh^/J; RRID:IMSR_JAX:013044). Mice were housed at 28°C on a reversed 12hr light-dark cycle with ad libitum access to food and water. Experiments were performed during the animals’ dark cycle and housed individually in an enriched environment after major surgery. Euthanasia was performed by infusion of a ketamine (300mg/kg) and dexmedetomidine (3mg/kg) or prolonged exposure to CO2. Both methods are consistent with the recommendations of the American Veterinary Medical Association (AVMA) Guidelines on Euthanasia.

### Surgical Procedures

Mice were induced to a surgical plane of anesthesia with 3% isoflurane in oxygen and secured in the nose cone of a stereotaxic frame. Animals were then maintained in an anesthetic plane with 1-2% isoflurane in oxygen and kept at a body temperature of 36° C with the use of a homeothermic blanket. Prior to any surgical procedure, mice were administered subcutaneous injections of sustained release buprenorphine (1 mg/kg) for analgesia, dexamethasone (0.2 mg/kg) for reduction of swelling, and bupivacaine (2 mg/kg), for local anesthesia. Mice received a subcutaneous injection of an antibiotic, enrofloxacin (Baytril; 5 mg/kg), for 3 days as part of their post-operative care.

### Viral Injections

Approximately 21 days prior to electrophysiological recordings or transcardial perfusions, we performed small (0.5 mm in diameter) unilateral craniotomies using a micromotor drill (Foredom)–under aseptic conditions–over auditory TRN (audTRN) or MGB. Glass syringes attached to a syringe pump (Pump 11 Elite, Harvard Apparatus) were backfilled with modified viral vectors, placed over the brain region of interest, and used to inject virus at 60 nL/min. Glass syringes were made using a micropipette puller (P-97, Sutter Instruments) from glass capillaries (Harvard Apparatus, 30-0038) with tip openings ranging from 30-40 µm in diameter. Following injections, syringes were left in place for 15-20 min before retraction. Craniotomies were then covered with bone wax (Fine Science Tools) or a removable silicone plug (Kwik-Cast, World Precision Instruments). The coordinates used to target audTRN were: -1.80 mm posterior to bregma, ±2.25 mm lateral to bregma, -2.90 mm below the pial surface. The MGB coordinates used were: -3.30 mm posterior to bregma, ±2.00 mm lateral to bregma, -2.90 mm below the pial surface.

### Viral Vectors

For soma-targeted inactivation of PV or SST neurons of the audTRN we injected 400 nL of AAV5-hSyn1-SIO-stGtACR1-FusionRed (≥7×10^12^ vg/mL, Addgene 105678-AAV5). For anterograde tracing of PV or SST neurons of the audTRN and control experiments we injected 400-450 nL of AAV5-hSyn-DIO-mCherry (≥ 7×10^12^ vg/mL, Addgene 50459-AAV5).

### Fiber Optic Cannula & Headpost Implantation

Following virus injections, fiber optic cannulas (Prizmatix: 3 mm in length, 1.25 mm in diameter, Ø 200 µm core, 0.66 NA) were implanted above audTRN at a 20° angle, -1.80 mm posterior to bregma, ±3.25 mm lateral to bregma, and -2.50 mm below the pial surface. A small craniotomy was made over the left frontal lobe and a ground pin was implanted. Craniotomies were sealed with Kwik-Cast (World Precision Instruments). The cannula, ground pin and a custom-made stainless-steel headpost (eMachine shop) were then secured with dental cement (C&B Metabond, Parkell) and dental acrylic (Lang Dental).

### Tissue Processing

After allowing time for proper viral expression or following electrophysiological recordings, mice were deeply anesthetized with a cocktail of ketamine and dexmedetomidine (*see Animals*) and transcardially perfused with 1X PBS followed by 4% paraformaldehyde (PFA) in PBS. Brains were extracted and post-fixed in 4% PFA overnight at 4° C. Following post-fixation, tissue was cryoprotected in 30% sucrose for at least 24 h at 4° C until sectioning. Brain sections 40-50 µm in thickness were cut on a cryostat (Leica CM1860) and collected for immunohistochemistry or imaging of viral expression. For imaging of viral expression or probe location, 50 µm sections were mounted on gelatin-coated glass slides and coverslipped using ProLong Diamond Antifade Mounting media with DAPI (Invitrogen P36971). For immunohistochemistry, 40 µm sections were collected in 12-well plates filled with 1X PBS (5-10 sections per well). Slides were imaged using a fluorescent microscope (Leica DM6 B) or a confocal microscope (Zeiss LSM 800).

### Immunohistochemistry

Free-floating 40 µm-thick brain sections were first washed in 1X PBS (3 x 10 min washes). To increase membrane permeability, sections were microwaved for approximately 10 s and then incubated in a solution of 0.3% Triton X-100 in 1X PBS (hereinafter referred to as PBT) for 30 min at room temperature. The sections were then incubated in a blocking solution consisting of 5% normal goat serum in PBT (hereinafter referred to as PBTG) for 0.5-1 h at room temperature. Following incubation with PBTG, sections were incubated in primary antibody solution which consisted of the following antibodies diluted in PBTG: (1) 1:500 mouse monoclonal anti-Parvalbumin (Swant, PV-235; RRID:AB_10000343), (2) 1:500 rabbit polyclonal anti-Somatostatin (Immunostar, 20067); RRID: AB_572264), and (3) 1:1000 rabbit polyclonal anti-Calretinin (Swant, Calretinin-7697; RRID:AB_2721226). Incubation in primary solution was done overnight-48 h at 4° C on a shaker plate (Fisher Scientific); well plates were covered with parafilm and a plastic lid. Following incubation with primary antibody solution, sections were washed in PBT (3 x 10 min washes). Sections were then incubated in secondary antibodies conjugated to fluorescent markers diluted in PBTG for 2 h at room temperature; well plates were covered with aluminum foil to block light. Secondary antibodies used were: (1) for PV staining, Alexa Fluor Plus 488 Goat anti-Mouse IgG (H+L) (Invitrogen A32723; RRID: AB_2633275), (2) for SST staining, Alexa Fluor Plus 488 Goat anti-Rat IgG (H+L) (Invitrogen A48262; RRID: AB_2896330), and (2) for Calretinin staining, Alexa Fluor Plus 488 Goat anti-Rabbit IgG (H+L) (Invitrogen A32731; RRID: AB_2633280). Sections were then incubated for 10 min in a 1:40,000 dilution of DAPI stock solution in PBT (5 mg/mL, Invitrogen D1306). The sections were subsequently washed in 1X PBS (5 x 10 min washes). Sections were then mounted on gelatin-coated glass slides, coverslipped using ProLong Diamond Antifade Mounting media (Invitrogen P36970) and left to curate in the dark, on a flat surface at room temperature for at least 24 h prior to imaging. Slides were imaged using a fluorescence microscope (Leica DM6 B) or a confocal microscope (Zeiss LSM 800).

### Acoustic Stimuli

Stimuli were generated using custom MATLAB code and were sampled at 200 kHz with 32-bit resolution. Acoustic stimuli were delivered through a speaker (Newark Multicomp, MCPCT-G5100-4139) positioned in the direction of the animals’ ear contralateral to the recording site. To test for stimulus-locked auditory responses, we used a set of click trains made up of 5 broadband noise bursts at a rate of 1 Hz. The noise bursts were 25 ms long and presented at a rate of 10 Hz for a total of 500 ms. To assess frequency response functions in neuronal populations, we generated a set of 19 pure tones of logarithmically spaced frequencies ranging between 3 kHz and 80 kHz at 70 dB sound pressure level relative to 20 μPa (SPL). Each tone was 50 ms long with a 1-ms cosine squared ramp, repeated 40 times with an inter-stimulus-interval of 300 ms and presented in a pseudo-random order. 20 of those 40 tone repetitions were accompanied by a continuous light pulse with a 1-ms cosine squared ramp starting at tone onset and ending at tone offset (50 ms; for details on light stimuli see Optogenetic Inactivation).

### Optogenetic Inactivation and Calibration

To inactivate populations of TRN neurons, we injected a soma-targeted inhibitory opsin in the audTRN of PV- and SST-Cre mice (see Viral Injections and Viral Vectors). On light-Only trials, we delivered continuous (1-ms cosine squared ramp) light pulses of different durations (10 ms, 25 ms, 50 ms, and 100 ms) through an implanted fiber-optic cannula via a fiber-coupled blue LED (460 nm, Prizmatix, Optogenetics-LED-Blue). During sound presentation, on light-On trials, we delivered a continuous 50-ms pulse with a 1-ms squared ramp concurrent with the tone stimulus.

To reduce variability in the effects of optogenetic manipulation across mice, we developed a procedure to calibrate the LED intensity prior to each recording session. Once the probe was in its final depth for recording, we presented alternating click trains (*see Acoustic Stimuli*) where each trial consisted of a click train without LED stimulation, followed by a click train paired with LED stimuli with one of 20 intensities (LED driving voltage ranged from 0 to 5 V) while recording neural responses. This stimulus set was presented 10 times for each intensity value. Neural responses were sorted offline using Kilosort2. To calculate if each unit was significantly sound responsive and discard noise clusters, we computed a Wilcoxon sign-rank test between sound responses during the light-Off click trains and a baseline period of activity. We excluded any units with P-values exceeding a Bonferroni-corrected α-level of 0.05/n_clusters_ from further analysis. We then calculated the number of spikes during the light-Off click train trials (n_OFF_) and during the light-On click train trials (n_ON_) and computed the percent change in the response (r_PC_) for each light intensity:

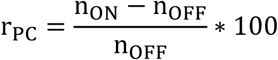

Using a logistic function, we fit the response function as a function of the LED voltage,

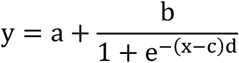

where a determines the y-offset of the response, b determines the range of the response, c determines the x-offset, and d determines the gain of the response. With this fit equation, we determined the voltage value driving the LED that elicited a 100% increase (2-fold change) in neural response and we used that voltage for all subsequent LED stimuli presented in each individual recording session (0.1-5 mW). Some sessions did not produce a 2-fold increase with LED manipulation; for these sessions we used maximum LED power (5 mW) for all stimuli.

### Acute Electrophysiological Recordings

All electrophysiological recordings were carried out in a double-walled acoustic-isolation booth (Industrial Acoustics). At least 21 days following viral injection, animals were anesthetized (see Surgical Procedures) and a small craniotomy (1 mm in diameter) was performed using a micromotor drill (Foredom) over MGB (for coordinates see Viral Injections). Mice were then clamped into a custom base and allowed to recover from anesthesia for at least 30 min. Following recovery, we vertically (0° on stereotaxic arm) lowered a 32-channel silicon probe (NeuroNexus: A1x32-Poly2-10mm-50s-177) to a depth of ∼3.5 mm from the pial surface using a motorized micromanipulator (Scientifica) attached to a stereotaxic arm (Kopf) at a rate of 1-2 µm/s. While we lowered the probe, a train of brief broadband noise clicks was played (*see Acoustic Stimuli*). If we observed stimulus-locked responses, we determined the probe was in a sound-responsive area. After some recordings, the probe was coated with a lipophilic far-red dye (Vybrant DiD, Invitrogen) and lowered in the recording location to determine recording sites (*see Tissue Processing*; **Figure 2C**). Recordings that did not display stimulus-locked responses post-hoc were not used for further analysis. Once the probe reached the target recording depth, it was left to settle in the brain for at least 30 min prior to recording. Neural signals were amplified via an Intan headstage (RHD 32ch, Intan Technologies), recorded at a rate of 30 kHz using an OpenEphys acquisition board and GUI^78^, and digitized with an SPI cable. Signals were filtered between 500 and 6000 Hz, offset corrected, and re-referenced to the median of all active channels. Recorded neural data were spike sorted using KiloSort2^79^ and manually corrected using phy2. Upon manual correction, units were classified as either single- or multi-units; if units exhibited a clear refractory period they were labeled single units, otherwise they were classified as multi-units. We later tested the proportions of single- and multi-units that were inhibited, facilitated or unchanged by SST and PV inactivation and found no differences between the groups (**Supplementary Figure S8**); therefore both single and multi-units were included in subsequent analyses (for number of units used see **Table 1**). Approximately 3 recording sessions were acquired from each individual mouse at different recording sites. For analysis in **Supplementary Figure S4**, we fit linear fit to recordings depth versus best frequency of neurons for each session, and designated sessions with a positive slope and significant p value (p< 0.05) as tonotopic, whereas other recording sites were designated as non-tonotopic ^80,81^.

**Table 1.** Statistics Table.

| Figure | Comparison | N | Mean | Median | SEM | Test | p |
| --- | --- | --- | --- | --- | --- | --- | --- |
| **1B.** | Gray Values; PV-audTRN projections; dMGB vs. vMGB | 14 | d:<br>30.18 | d:<br>26.33 | d:<br>2.65 | Wilcoxon signed-rank test | 0.000 |
|  |  |  | v:<br>41.41 | v:<br>39.53 | v:<br>2.65 |  |  |
| **1D.** | Gray Values; SST-audTRN projections; dMGB vs. vMGB | 8 | d:<br>95.04 | d:<br>102.71 | d:<br>17.17 | Wilcoxon signed-rank test | 0.023 |
|  |  |  | v:<br>58.06 | v:<br>62.84 | v:<br>5.83 |  |  |
| **2** | $\Delta$ FR (Hz); PV-facilitated.<br>2.5-2.9 mm (1<sup>st</sup> bin) vs. -3.1-3.5 mm (2<sup>nd</sup> bin) | 1<sup>st</sup> bin:<br>16 | 1<sup>st</sup> bin:<br>12.45 | 1<sup>st</sup> bin:<br>5.61 | 1<sup>st</sup> bin:<br>3.72 | Mann-Whitney U Test | 0.008 |
|  |  | 2<sup>nd</sup> bin:<br>30 | 2<sup>nd</sup> bin:<br>41.66 | 2<sup>nd</sup> bin:<br>24.70 | 2<sup>nd</sup> bin:<br>11.16 |  |  |
| **2** | $\Delta$ FR (Hz); PV-Suppressed.<br>2.5-2.9 mm (1<sup>st</sup> bin) vs. -3.1-3.5 mm (2<sup>nd</sup> bin) | 1<sup>st</sup> bin:<br>30 | 1<sup>st</sup> bin:<br>-10.35 | 1<sup>st</sup> bin:<br>-9.60 | 1<sup>st</sup> bin:<br>0.97 | Mann-Whitney U Test | 0.219 |
|  |  | 2<sup>nd</sup> bin:<br>39 | 2<sup>nd</sup> bin:<br>-15.31 | 2<sup>nd</sup> bin:<br>-12.28 | 2<sup>nd</sup> bin:<br>2.14 |  |  |
| **2** | $\Delta$ FR (Hz); SST-facilitated.<br>2.5-2.9 mm (1<sup>st</sup> bin) vs. -3.1-3.5 mm (2<sup>nd</sup> bin) | 1<sup>st</sup> bin: 0 | 1<sup>st</sup> bin:<br>n/a | 1<sup>st</sup> bin:<br>n/a | 1<sup>st</sup> bin:<br>n/a | n/a | n/a |
|  |  | 2<sup>nd</sup> bin: 2 | 2<sup>nd</sup> bin:<br>6.57 | 2<sup>nd</sup> bin:<br>6.57 | 2<sup>nd</sup> bin:<br>0.48 |  |  |
| **2** | $\Delta$ FR (Hz); SST-Suppressed;<br>2.5-2.9 mm (1<sup>st</sup> bin) vs. -3.1-3.5 mm (2<sup>nd</sup> bin) | 1<sup>st</sup> bin:<br>51 | 1<sup>st</sup> bin:<br>-13.46 | 1<sup>st</sup> bin:<br>-10.85 | 1<sup>st</sup> bin:<br>1.29 | Mann-Whitney U Test | 0.012 |
|  |  | 2<sup>nd</sup> bin:<br>66 | 2<sup>nd</sup> bin:<br>-10.37 | 2<sup>nd</sup> bin:<br>-7.08 | 2<sup>nd</sup> bin:<br>1.3 |  |  |
| **3** | Light responses (Hz), PV-facilitated, light Only vs. light diff (tone+light – tone only) vs, 0-25 ms, | 74 | Light Only:<br>29.19 | Light Only:<br>11.45 | Light Only:<br>6.35 | Wilcoxon signed-rank test t | 0.0113 |
|  |  |  | Light diff:<br>33.47 | Light diff:<br>13.37 | Light diff:<br>6.35 |  |  |
| **3** | Light responses (Hz), PV-suppressed, light Only vs. light diff (tone+light – tone only) vs, 0-25 ms | 104 | Light-Only:<br>- 2.75 | Light-Only:<br>- 2.53 | Light-Only:<br>0.66 | Wilcoxon signed-rank test t | 1.16e-12 |
|  |  |  | Light diff:<br>-12.59 | Light diff:<br>-10.10 | Light diff:<br>1.02 |  |  |
| **3** | Time to peak (ms); PV-Facilitated: Light<sub>off</sub> vs. Light<sub>on</sub> | 74 | Off:<br>12.27 | Off:<br>12.0 | Off:<br>0.70 | Wilcoxon signed-rank test | 0.0398 |
|  |  |  | On:<br>13.7 | On:<br>14.0 | On:<br>0.56 |  |  |
| **3** |  | 104 | Off:<br>9.19 | Off:<br>8 | Off:<br>0.51 |  | 0.070 |
|  | Time to peak (ms); PV-Suppressed: Light<sub>Off</sub> vs. Light<sub>On</sub> |  | On:<br>8.73 | On:<br>8 | On:<br>0.53 | Wilcoxon signed-rank test |  |
| 3 | Firing Rate (Hz)<br>0-25 ms; PV Facilitated: Light<sub>Off</sub> vs. Light<sub>On</sub> | 74 | Off:<br>14.58 | Off:<br>7.20 | Off:<br>2.118 | Wilcoxon signed-rank test | 0.00000 |
|  |  |  | On:<br>48.05 | On:<br>28.78 | On:<br>6.85 |  |  |
| 3 | Firing Rate (Hz)<br>0-25 ms; PV-Suppressed: Light<sub>Off</sub> vs. Light<sub>On</sub> | 104 | Off:<br>24.34 | Off:<br>17.71 | Off:<br>1.93 | Wilcoxon signed-rank test | 0.00000 |
|  |  |  | On:<br>11.76 | On:<br>5.02 | On:<br>1.37 |  |  |
| 3 | Firing Rate (Hz)<br>25-50 ms; PV Facilitated: Light<sub>Off</sub> vs. Light<sub>On</sub> | 74 | Off:<br>22.45 | Off:<br>14.57 | Off:<br>2.96 | Wilcoxon signed-rank test | 0.00004 |
|  |  |  | On:<br>36.40 | On:<br>24.72 | On:<br>4.97 |  |  |
| 3 | Firing Rate (Hz)<br>25-50 ms; PV Suppressed: Light<sub>Off</sub> vs. Light<sub>On</sub> | 104 | Off:<br>14.96 | Off:<br>7.42 | Off:<br>2.67 | Wilcoxon signed-rank test | 0.00000 |
|  |  |  | On:<br>4.88 | On:<br>1.27 | On:<br>1.13 |  |  |
| 3 | Light responses (Hz), PV-facilitated, light Only vs. light diff (tone+light – tone only) vs, 0-25 ms, | 74 | Light Only:<br>29.19 | Light Only:<br>11.45 | Light Only:<br>6.35 | Wilcoxon signed-rank test t | 0.0113 |
|  |  |  | Light diff:<br>33.47 | Light diff:<br>13.37 | Light diff:<br>6.35 |  |  |
| 3 | Light responses (Hz), PV-suppressed, light Only vs. light diff (tone+light – tone only) vs, 0-25 ms | 104 | Light-Only:<br>- 2.75 | Light-Only:<br>- 2.53 | Light-Only:<br>0.66 | Wilcoxon signed-rank test t | 1.16e-12 |
|  |  |  | Light diff:<br>-12.59 | Light diff:<br>-10.10 | Light diff:<br>1.02 |  |  |
| 4 | Time to peak (ms); SST Facilitated: Light<sub>Off</sub> vs. Light<sub>On</sub> | 4 | Off:<br>12.0 | Off:<br>11 | Off:<br>2.16 | Wilcoxon signed-rank test | 1 |
|  |  |  | On:<br>12.0 | On:<br>12 | On:<br>2.58 |  |  |
| 4 | Time to peak (ms); SST Suppressed: Light<sub>Off</sub> vs. Light<sub>On</sub> | 181 | Off:<br>9.23 | Off:<br>8 | Off:<br>0.378 | Wilcoxon signed-rank test | 0.071 |
|  |  |  | On:<br>8.95 | On:<br>6 | On:<br>0.421 |  |  |
| 4 | Firing Rate (Hz), 0-25 ms; SST Facilitated: Light<sub>Off</sub> vs. Light<sub>On</sub> | 4 | Off:<br>30.31 | Off:<br>32.41 | Off:<br>8.42 | Wilcoxon signed-rank test | 0.125 |
|  |  |  | On:<br>36.34 | On:<br>38.74 | On:<br>8.30 |  |  |
| 4 | Firing Rate (Hz), 0-25 ms; SST Suppressed: Light<sub>Off</sub> vs. Light<sub>On</sub> | 181 | Off:<br>24.43 | Off:<br>16.19 | Off:<br>1.73 | Wilcoxon signed-rank test | 0.00000000 |
|  |  |  | On:<br>12.86 | On:<br>6.19 | On:<br>1.24 |  |  |
| 4 | Firing Rate (Hz) 25-50 ms; SST Facilitated: Light<sub>Off</sub> vs. Light<sub>On</sub> | 4 | Off:<br>8.51 | Off:<br>5.07 | Off:<br>4.63 | Wilcoxon signed-rank test | 0.125 |
|  |  |  | On:<br>12.86 | On:<br>6.19 | On:<br>1.24 |  |  |
| 4 | Firing Rate (Hz) 25-50 ms; SST Suppressed: Light<sub>Off</sub> vs. Light<sub>On</sub> | 181 | Off:<br>10.59 | Off:<br>5.22 | Off:<br>1.22 | Wilcoxon signed-rank test | 0.00000 |
|  |  |  | On:<br>7.39 | On:<br>2.23 | On:<br>1.05 |  |  |
| 4 | Light responses (Hz), SST-Facilitated, light Only vs. light diff (tone+light – tone only) vs, 0-25 ms, | 4 | Light Only: -4.23 | Light Only: -2.49 | Light Only: 2.74 | Wilcoxon signed-rank test | 0.026 |
|  |  |  | Light diff: 6.038 | Light diff: 5.86 | Light diff: 0.37 |  |  |
| 4 | Light responses (Hz), SST-Suppressed, light Only vs. light diff (tone+light – tone only) vs, 0-25 ms, | 181 | Light Only: -4.21 | Light Only: -2.17 | Light Only: 0.47 | Wilcoxon signed-rank test | 3.8e-18 |
|  |  |  | Light diff: -11.57 | Light diff: -7.91 | Light diff: 0.80 |  |  |
| 5 | Sparseness, 0-25 ms; PV Facilitated. Light<sub>Off</sub> vs. Light<sub>On</sub> | 74 | Off: 0.476 | Off: 0.487 | Off: 0.030 | Wilcoxon signed-rank test | 0.000 |
|  |  |  | On: 0.179 | On: 0.122 | On: 0.020 |  |  |
| 5 | Sparseness, 0-25 ms; PV Suppressed. Light<sub>Off</sub> vs. Light<sub>On</sub> | 104 | Off: 0.383 | Off: 0.356 | Off: 0.017 | Wilcoxon signed-rank test | 0.00000009 |
|  |  |  | On: 0.468 | On: 0.454 | On: 0.020 |  |  |
| 5 | Sparseness, 0-25 ms; SST Facilitated. Light<sub>Off</sub> vs. Light<sub>On</sub> | 4 | Off: 0.249 | Off: 0.273 | Off: 0.044 | Wilcoxon signed-rank test | 0.125 |
|  |  |  | On: 0.167 | On: 0.159 | On: 0.028 |  |  |
| 5 | Sparseness, 0-25 ms; SST Suppressed. Light<sub>Off</sub> vs. Light<sub>On</sub> | 181 | Off: 0.361 | Off: 0.332 | Off: 0.013 | Wilcoxon signed-rank test | 0.00000000 |
|  |  |  | On: 0.518 | On: 0.518 | On: 0.015 |  |  |
| S6 | Mu (change, Hz), PV Suppressed. Light<sub>Off</sub> vs. Light<sub>On</sub> | 26 | -3159.1 | 209.2 |  | Wilcoxon signed-rank test | 1 |
| S6 | Mu (change, Hz). PV Facilitated. Light<sub>Off</sub> vs. Light<sub>On</sub> | 11 | -1308.2 | -497.1 |  | Wilcoxon signed-rank test | 0.465 |
| S6 | Sigma (change). PV Facilitated. Light<sub>Off</sub> vs. Light<sub>On</sub> | 11 | 0.068 | .0.03 |  | Wilcoxon signed-rank test | 0.278 |
| S6 | Sigma (change). PV Suppressed. Light<sub>Off</sub> vs. Light<sub>On</sub> | 26 | -0.124 | -0.072 |  | Wilcoxon signed-rank test | 0.027 |
| S6 | Mu (change, Hz). SST Facilitated. Light<sub>Off</sub> vs. Light<sub>On</sub> | 0 | n/a | n/a | n/a |  |  |
| S6 | Mu (change, Hz). SST Suppressed. Light<sub>Off</sub> vs. Light<sub>On</sub> | 47 | 2672.5 | 156.1 |  | Wilcoxon signed-rank test | 0.021 |
| **S6** | Sigma (change). SST Facilitated. Light<sub>Off</sub> vs. Light<sub>On</sub> | 0 | n/a | n/a | n/a |  |  |
| **S6** | Sigma (change). SST Suppressed. Light<sub>Off</sub> vs. Light<sub>On</sub> | 47 | -0.19 | -0.06 |  | Wilcoxon signed-rank test | 0.0002 |
| **S9** | Firing Rate (Hz), 0-25 ms; PV-Cre, mCherry (no opsin): Light<sub>Off</sub> vs. Light<sub>On</sub> | 85 | Off: 14.76 | Off: 8.10 | Off: 2.31 | Wilcoxon signed-rank test | 0.087 |
|  |  |  | On: 15.08 | On: 8.70 | On: 2.30 |  |  |
| **S9** | Firing Rate (Hz) 25-50 ms; PV-Cre, mCherry (no opsin): Light<sub>Off</sub> vs. Light<sub>On</sub> | 85 | Off: 6.98 | Off: 3.73 | Off: 1.08 | Wilcoxon signed-rank test | 0.762 |
|  |  |  | On: 9.50 | On: 3.92 | On: 1.72 |  |  |
| **S9** | Firing Rate (Hz) 50-75 ms; PV-Cre, mCherry (no opsin): Light<sub>Off</sub> vs. Light<sub>On</sub> | 85 | Off: 5.79 | Off: 4.16 | Off: 0.75 | Wilcoxon signed-rank test | 0.434 |
|  |  |  | On: 7.05 | On: 4.02 | On: 0.91 |  |  |
| **S10** | Firing Rate (Hz), 0-25 ms; SST-Cre, mCherry (no opsin): Light<sub>Off</sub> vs. Light<sub>On</sub> | 137 | Off: 16.19 | Off: 12.61 | Off: 1.24 | Wilcoxon signed-rank test | 0.373 |
|  |  |  | On: 16.38 | On: 13.38 | On: 1.25 |  |  |
| **S10** | Firing Rate (Hz) 25-50 ms; SST-Cre, mCherry (no opsin): Light<sub>Off</sub> vs. Light<sub>On</sub> | 137 | Off: 12.40 | Off: 8.53 | Off: 1.08 | Wilcoxon signed-rank test | 0.006 |
|  |  |  | On: 15.07 | On: 10.53 | On: 1.36 |  |  |
| **S10** | Firing Rate (Hz) 50-75 ms; SST-Cre, mCherry (no opsin): Light<sub>Off</sub> vs. Light<sub>On</sub> | 137 | Off: 9.99 | Off: 7.95 | Off: 0.72 | Wilcoxon signed-rank test | 0.000002 |
|  |  |  | On: 12.25 | On: 8.92 | On: 0.95 |  |  |

### Gray Value Analysis & Cell Counting

*Gray Value:* We first delineated the MGB across the different ROIs. We then used the Fiji Plot Profile plugin to quantify gray values. We used a custom Python script to calculate the mean gray values and S.E.M. *Cell Counting:* We used the Cellpose^82^ anatomical segmentation algorithm to outline cells in selected sections. We then manually confirmed the masks provided by Cellpose in Fiji using the cell counter plugin. A custom Python script was used to calculate the overlapping masks and percentages.

### Neural Response Analysis

We selected units for further analysis for inclusion based on their stimulus response profiles. We repeated experiments after injecting a virus encoding only the fluorescent reporter without an opsin and found no significant effect of light on neuronal responses 0-25 ms after tone onset, but a mixed effect of light on responses 25-50 ms and 50-75 ms after tone onset: no effect for PV-Cre mice, but a significant effect in SST-Cre mice (**Supplementary Figures S9, S10**). This effect likely corresponds to delayed visual responses and may be due to leakage of light between the skull and the fiberoptic cannula. Therefore, we focused our main analyses on responses 0-25 ms after tone onset. To select sound- and light-responsive units, we fit an ordinary least square linear regression model to mean responses on each trial during 0-25 ms after the sound onset with sound and light as predictors and a (light x sound) interaction predictor. We selected units that had significant sound predictor as sound-responsive; and units that were significant for light responses as those with significant light or light-sound interaction predictors (α < 0.05). Spontaneous firing rate was computed as the average firing rate during the 25 ms prior to tone onset for light-Off and light-On trials separately. Tone-evoked firing rate was calculated as the average firing rate for the first 25 ms of the total tone duration for light-Off and light-On trials separately. For some figures, we also computed the firing rates in the late (25-50 ms) and offset (50-75 ms) periods after tone onset. Light-Only responses were calculated as the average firing rate for the first 25 ms of light-Only trials. We calculated differences between responses in light-On and light-Off conditions by subtracting the mean of the firing rates during the first 25 ms of tone presentation in the light-On trials from that in light-Off trials: light-responsive cells with positive difference were classified as facilitated, and with negative difference as suppressed. To calculate frequency response functions, we averaged the firing rate during the first 25 ms of tone presentation for each trial of the 19 frequencies tested in both light-Off and light-On conditions. Best frequency was defined as the frequency that elicited the largest change in firing rate compared to baseline activity. Sparseness for each neuron was calculated separately for the light-Off and light-On conditions during the first 25 ms of tone presentation. The sparseness equation used is as follows:

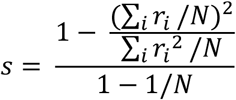

where r_i_ is the firing rate of a single neuron at each frequency and N is the number of frequencies tested. To examine how frequency tuning changes in light-On and light-Off conditions (**Supplementary Figure S6**), we fit a Gaussian curve of the form

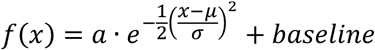

to the mean frequency response function of each unit. We initialized each fit four times using different initial parameters, calculated R^2^, and selected the fit that explained the highest amount of variance in the data (highest R^2^).

### Statistical Analysis

We used ordinary least squares regression with light and tone as factors to assay for significant effect on individual neuronal responses over trials, as well as for significance of their interactions using ols function in Python. We assessed normality of the data using a Shapiro-Wilk tests using Scipy’s Stats Python Library. For paired data that violated the assumption of normality, p values were calculated using a Wilcoxon sign-rank tests using Scipy’s Stats Python Library. For non-paired data that violated the assumption of normality, p values were calculated using a Wilcoxon rank-sum tests using Scipy’s Stats Python Library. The standard error of the mean was used to calculate error bars (± SEM). Symbols: * indicates p values <0.05, ** indicates p values < 0.01, and *** indicates p values <0.001.

### Computational Modeling

Model cells were Hodgkin-Huxley formalism solved by a second order Runge-Kutta ODE solver in MATLAB (ode23, Mathworks). TRN and MGB cells were single-compartment Hodgkin-Huxley models built upon those used previously ^48,83,84^ that include T currents. To match the reduced bursting properties in SST^TRN^ and higher-order MGB cells^43,85^, those cells had 50% reduced T current densities (0.375 mS/cm^2^). Parameters included the following ionic currents and maximal conductances: fast transient Na+ (NaT) 60.5 mS/cm^2^, K+ delayed rectifier (Kd) 60 mS/cm^2^, K+ transient A (Kt) 5 mS/cm^2^, slowly inactivating K+ (K2) 0.5 mS/cm^2^, slow anomalous rectifier (AR) 0.025 mS/cm^2^, and low threshold transient Ca2+ (CaT) 0.75 mS/cm^2^ for PV/MGB cells and 0.375 mS/cm^2^ for SST/HO cells. Reversal potentials were 50 mV for sodium, -100 mV for potassium, 125 mV for calcium, -40 mV for AR and -75 mV for leak. Capacitance was 1 µF/cm^2^ with leak of 0.1 mS/cm^2^. Chemical synapses were modelled as double exponential decay with rise and fall time kinetics of 5 ms and 35 ms respectively, with reversal potentials of 0 mV for excitatory and -100 mV for inhibitory synapses. Electrical synapses were modelled as static coupling conductance of 0.03 mS/cm^2^ applied to the voltage difference between the coupled TRN cells, corresponding to a strong electrical coupling (∼0.2 coupling coefficient). Tone inputs were simulated as exponentially decaying current pulses (peak amplitude 2.7 µA/cm^2^, with decay time constant 30 ms) delivered simultaneously to each of the MGB cells. Each of the TRN cells was inactivated individually through a static conductance of 10 mS/cm^2^ and reversal potential of -70 mV for 100 ms during the inputs (**Figure 7B**). Chemical synaptic connections between TRN cells were set at 1 µA/cm^2^ for excitatory synapses from MGB to TRN cells, and 3 µA/cm^2^ for all inhibitory synapses originating from TRN. For each stimulus condition, we repeated 150 trials under Poisson random noise of 0.2 µA at a rate of 80 Hz for excitatory and 20 Hz for inhibitory events. Among these trials, we varied the percent of trials that included various synaptic connections from TRN cells (**Figure 7C**), in lieu of varying synaptic strengths. Subthreshold current steps of 0.3 µA were applied to all cells to increase excitability. PSTHs were obtained from spike time data histograms with a bin size of 1 ms and smoothed with a 31 ms Hanning window. We normalized each response to the maximum rate in the non-inactivated trial.

## RESOURCE AVAILABILITY

### Lead Contact

Further information and requests for resources and reagents should be directed to and will be fulfilled by the Lead Contact, Maria Geffen.

### Data & Code availability

Source data has been deposited in Dryad: https://doi.org/10.5061/dryad.ht76hdrq5. Data analysis code is available on GitHub: https://github.com/geffenlab/Rolon_Martinez_2024. Code for the model is available at github: https://github.com/jhaaslab/TRN_MGB_celltypespecific.

## Acknowledgements

The authors would like to thank the members of the Geffen and Haas laboratories and Yale Cohen for their discussions and helpful comments.

## Financial Disclosure

This work was supported by the following grants from NIH: R01DC15527, R01DC014479, R03DC013660 to MNG, F31DC018473 to SRM, F31DC016524to CFA, from NINDS R01NS113241 to MNG and R01 NS128713 to JSH. The funders had no role in study design, data collection and analysis, decision to publish, or preparation of the manuscript.

**Supplementary Figure Legends.**

**Supplementary Figure S1. Co-expression of PV+ and SST+ neurons with the inhibitory soma targeted opsin stGtACR1.FusionRed. A.** *Left*: AAV5.hSyn.SIO.stGtACR1.FusionRed expression in the audTRN of a PV-Cre mouse. *Middle*: Immunohistochemical expression of parvalbumin-positive neurons. *Right*: Merged image of PV-audTRN neurons expressing both the parvalbumin antibody and the inhibitory opsin. **B**. Quantification of PV+ neurons, PV+stGtACR1.FusionRed+ neurons, and stGtACR1.FusionRed+ neurons. **C.** *Left*: AAV5.hSyn.SIO.stGtACR1.FusionRed expression in the audTRN of an SST-Cre mouse. *Middle*: Immunohistochemical expression of somatostatin-positive neurons. *Right*: Merged image of SST-audTRN neurons expressing both the somatostatin antibody and the inhibitory opsin. **D**. Quantification of SST+ neurons, SST+stGtACR1.FusionRed+ neurons, and stGtACR1.FusionRed+ neurons. *Scale bar = 100µm. Arrows represent example cells. N=2 mice for each group*.

**Supplementary Figure S2. Optical inactivation of SST^TRN^ neurons suppresses tone-evoked activity in the TRN. A.** Average firing rate of recorded units in the TRN in response to 50 ms tone, with vs without optical inactivation of SST neurons (AAV5-hSyn1-SIO-stGtACR1-FusionRed in SST-cre mouse). n= 27 units. Shaded areas represent SEM±. **B.** Average firing rates of TRN units during baseline (Spont), tone-evoked responses (Tone) and offset responses (Offset), with vs without optical inactivation of SST neurons. ***p<0.001, Wilcoxon signed-rank tests. Error bars represent SEM±. **C.** Average firing rate (FR) of TRN units in response to tones on light ON versus light Off trials (with and without optical inactivation of SST neurons). ***p<0.001, Wilcoxon signed-rank test. **E.** Average frequency response function of TRN units with vs without optical inactivation of SST neurons. ***p<0.001, **p<0.01, *p<0.05, Wilcoxon signed-rank tests

**Supplementary Figure S3. A.** Overlay of live IR and fluorescence images of slice from PV-Cre mouse injected with AAV5_hSyn1-SIO-stGtACR1-FusionRed. **B.** Whole-cell recordings from a TRN neuron from (A) expressing GtACR1, injected with varying amplitudes of current (bottom) and exposed to widefield LED stimulation (blue bar). V^m^ = -80 mV. Scale bar 25 mV, 250 ms. *Methods for Supplementary Figure S3: 400* nL of AAV5_hSyn1-SIO-stGtACR1-FusionRed was injected to TRN (-2 AP, 1.35 ML, 3.0 and 3.2 depth; Kopf) of PV-Cre (Jax: 017320) animals under sterile anesthetized surgery. Mice were transcardially perfused with NMDG ACSF solution (in mM): 92 NMDG, 2.5 KCl, 1.25 NaH_2_PO_4_, 30 NaHCO_3_, 20 HEPES, 25 glucose, 2 thiourea, 5 NA ascorbate, 3 NA pyruvate, 0.5 CaCl_2_, and 10 MgSO_4_·7H_2_O (315 mOsm L−1, 7.3-7.4 pH). Horizontal brain slices 250 µm thick were cut and incubated in NMDG solution. Slices were incubated at 34°C for 12-15 min in NMDG following cutting and returned to HEPES ACSF at room temperature until recording. The HEPES ACSF solution comprised in mM: 92 NaCl, 2.5 KCl, 1.25 NaH_2_PO_4_, 30 NaHCO_3_, 20 HEPES, 25 glucose, 2 Thiourea, Na ascorbate, NA pyruvate, 2 CaCl_2_, and 2 MgSO_4_·7H_2_O (315 mOsm L−1, 7.3-7.4 pH). The ACSF bath during recording contained (in mM): 126 NaCl, 3 KCl, 1.25 NaH_2_PO_4_, 2 MgSO_4_, 26 NaHCO_3_, 10 dextrose, and 2 CaCl_2_ (315–320 mOsm L−1, saturated with 95% O2/5% CO_2_). The submersion recording chamber was held at 34°C (TC-324B, Warner Instruments). Electrodes were filled with (in mM): 135 potassium gluconate, 2 KCl, 4 NaCl, 10 Hepes, 0.2 EGTA, 4 ATP-Mg, 0.3 GTP-Tris, and 10 phosphocreatine-Tris (pH 7.25, 295 mOsm L^−1^). 1 M KOH was used to adjust pH of the internal solution. The approximate bath flowrate was 2 ml min^−1^ and the recording chamber held approximately 5 ml solution. TRN was visualized under 4x magnification, and TRN cells from auditory sector were identified and patched under 40x IR-DIC optics (SliceScope, Scientifica, Uckfield, UK). GtACR was excited by a 472 nm diode delivered through the objective (CoolLED pE-300). Voltage signals were amplified and low-pass filtered at 8 kHz (MultiClamp, Axon Instruments, Molecular Devices, Sunnyvale, CA,USA), digitized at 20 kHz with custom Matlab routines controlling a National Instruments (Austin, TX, USA, USB6221 DAQ board), and data were stored for offline analysis in Matlab (Mathworks, R2018b, Natick, MA, USA). Recordings were made in whole-cell current-clamp mode. Pipette resistances were 5-9 MΩ before bridge balance; recordings were discarded if access resistance exceeded 25 MΩ.

**Supplementary Figure S4. Similar tone and optogenetic responses for recorded MGB neurons in tonotopic and non-tonotopic recording sites in PV-cre mice.** A. Plot of neurons’ best frequency vs. recording depth for tonotopic recording sites; N= 47 neurons, 2 recording sites, 2 mice. Red line shows linear regression; r = .41, p = .0047. **B.** Pie chart breaking down the effect of PV^TRN^ inactivation in tonotopic recordings. N= 14 suppressed neurons (49%), 23 facilitated neurons (30%), 10 non-significant neurons (21%). **C, D.** Mean PSTH of facilitated (C) and suppressed (D) recorded neurons in MGB. Light-only trials (green line), tone-only trials (black line), and tone- and laser-on trials (light blue line). Shaded areas represent SEM±. **E-H.** Same as A-F, but for non-tonotopic recording sites. **G.** N= 209 neurons, 10 recording sites, 3 mice; r=-0.06, p=0.39. **H.** N= 81 suppressed neurons (39%), 60 facilitated neurons (29%), 68 non-significant neurons (33%).

**Supplementary Figure S5. Greater proportion of light-modulated MGB neurons in non-tonotopic than tonotopic recording sites during SST inactivation.** A. Plot of neurons’ best frequency vs recording depth for tonotopic recording sites; N= 39 neurons, 1 recording sites, 1 mouse. Red line shows linear regression; r = .67, p = 3e-6. **B.** Pie chart breaking down the effect of SST^TRN^ inactivation in tonotopic recordings. N= 2 facilitated neurons (5%), 8 suppressed neurons (21%), 29 non-significant neurons (74%). **C,D.** Mean PSTH of facilitated (E) and suppressed (F) recorded neurons in MGB. Light-only trials (green line), tone-only trials (black line), and tone- and laser-on trials (light blue line). Shaded areas represent SEM±. **E-H.** Same as A-F, but for non-tonotopic recording sites. **G.** N= 273 neurons, 9 recording sites, 4 mice; r=0.062, p=0.31. **H.** N= 173 suppressed neurons (63%), 2 facilitated neurons (1%), 98 non-significant neurons (36%).

**Supplementary Figure S6. Inactivation of PV^TRN^ and SST^TRN^ neurons shapes frequency tuning in MGB neurons. A.** scatterplot of μ for light-On and light-Off conditions for neurons facilitated by PV^TRN^ inactivation (first row) and neurons suppressed by PV^TRN^ inactivation (second row). **B.** Scatterplots of σ for light-On and light-Off conditions. **C.** Mean frequency response function for light-On (light blue) and light-Off (light gray) trials for neurons centered on μ (0 octaves = μ). **D.** A sample tuning curve for light-On (light blue) and light-Off (grey) conditions along with the gaussian fit for each curve. E-H. Sams as A-D, for suppressed MGB neurons during PV^TRN^ inactivation. **I-L.** Same as *A-D* for SST^TRN^ inactivation, suppressed MGB neurons. Facilitated neurons not show because only N = 1 neuronal responses were well-fitted by a Gaussian for both light on and light off conditions.

We first evaluated how well each tuning curve fit a Gaussian function in both the light-On and light-Off conditions (see Methods). We found that overall the number of MGB neurons with good Gaussian fits (R^2^ > 0.6 and µ/σ within tested range) was greater in the light-Off condition (n = 33/74 facilitated units and 58/104 suppressed units during PV^TRN^ inactivation; n = 84/181 suppressed units during SST^TRN^ inactivation) compared to the light-On condition (19/74 facilitated units and 43/104 suppressed units during PV^TRN^ inactivation; n = 88/181 suppressed neurons during SST^TRN^ inactivation) in all groups, except for neurons facilitated by SST^TRN^ activation (light-Off, n = 1/4 neurons, light-On, n = 1/4 neurons).

We found no differences in μ (best frequency) for neurons that had good fits in both light-On and light-Off conditions. We did not find significant differences between groups (units facilitated during PV^TRN^ inactivation: N = 11, t = 0.94, p = 0.37; units suppressed during PV^TRN^ inactivation: N = 26, t = 1.36, p = 0.186; units suppressed during SST^TRN^ inactivation: N = 47, t = -1.47, p = 0.147). MGB neurons that were suppressed by PV^TRN^ inactivation showed a significant decrease in σ (PV: N = 26, t = 2.42, p = 0.027; SST: N = 47, t = 3.30, p = 0.0002). However, for neurons facilitated by PV^TRN^ inactivation, there was no change in σ between light-On and light-Off conditions (N = 11, t = -1.27, p = 0.278) and there were no well-fit facilitated neurons during SST^TRN^ inactivation.

**Supplementary Figure S7. Optogenetic identification of TRN^SST^ to TRN^nonSST^ inhibitory synapse. A.** Widefield image of brain slice containing TRN from an SST-Cre mouse injected with AAV9-DIO-stChrome-mRuby. **B.** 40x image from A with patched neuron; no mRuby expression in this field. **C.** mRuby expression in another field from A confirming neuronal expression. **D.** Inward responses (IPSCs) from neuron patched with high-Cl^-^ internal solution following widefield optogenetic excitation (red bar). Scale bar 30 pA, 10 ms. Traces are offset for clarity. **E.** Peak-aligned average of responses from D. Scale bar 10 pA, 10 ms. Decay of the IPSC was best fit by a dual exponential with time constants 3.9 and 275 ms. **F.** Latencies of IPSCs from D.

**Supplementary Figure S8. Proportion of facilitated and suppressed MGB single units and multi-units. A.** Proportion of MGB MUA clusters (left) and single units (right) that were facilitated, suppressed, and non-significantly modulated by PV^TRN^ inactivation. **B**. Proportion of MGB MUA clusters (left) and single units (right) that were facilitated, suppressed, and non-significantly modulated by SST^TRN^ inactivation.

**Supplementary Figure S9. Control experiments for optogenetic manipulation of PV^TRN^ neurons. A.** PV mice were injected with an AAV5-hSyn1-DIO-mCherry virus into the audTRN and implanted with an optic fiber. We recorded from MGB neurons in awake head-fixed mice. We presented a set of pure tones in light-on and light-off conditions. **B**. We presented light in the audTRN of PV mice while recording from neurons in the MGB using a vertical multi-channel electrode that spanned the depth of dMGB and vMGB. **C.** Mean PSTH of recorded neurons in MGB. Light-only trials (green line), tone-only trials (black line), and tone- and laser-on trials (light blue line). **D**. Scatter plot of the mean FR for laser off and laser on conditions 0-25 ms (left), 25-50 ms (center), 50-75 ms (right) post-stimulus onset. *Shaded areas represent SEM±. N=85, n = 2; n.s. for all periods*.

**Supplementary Figure S10. Control experiments for optogenetic manipulation of SST^TRN^ neurons. A.** SST mice were injected with an AAV5-hSyn1-DIO-mCherry virus into the audTRN and implanted with an optic fiber. We recorded from MGB neurons in awake head-fixed mice. We presented a set of pure tones in light-on and light-off conditions. **B**. We presented light in the audTRN of SST mice while recording from neurons in the MGB using a vertical multi-channel electrode that spanned the depth of dMGB and vMGB. **C.** Mean PSTH of recorded neurons in MGB. Light-only trials (green line), tone-only trials (black line), and tone- and laser-on trials (light blue line). **D**. Scatter plot of the mean FR for laser off and laser on conditions 0-25 ms (left), 25-50 ms (center), 50-75 ms (right) post-stimulus onset. *Shaded areas represent SEM±. N=137, n = 2; p-value = n.s. (0-25 ms); 0.006 (25-50 ms), p = 0.000002 (50-75 ms)*.

