## Supplementary Figures for "Cell type-specific inhibitory modulation of sound processing in the auditory thalamus"

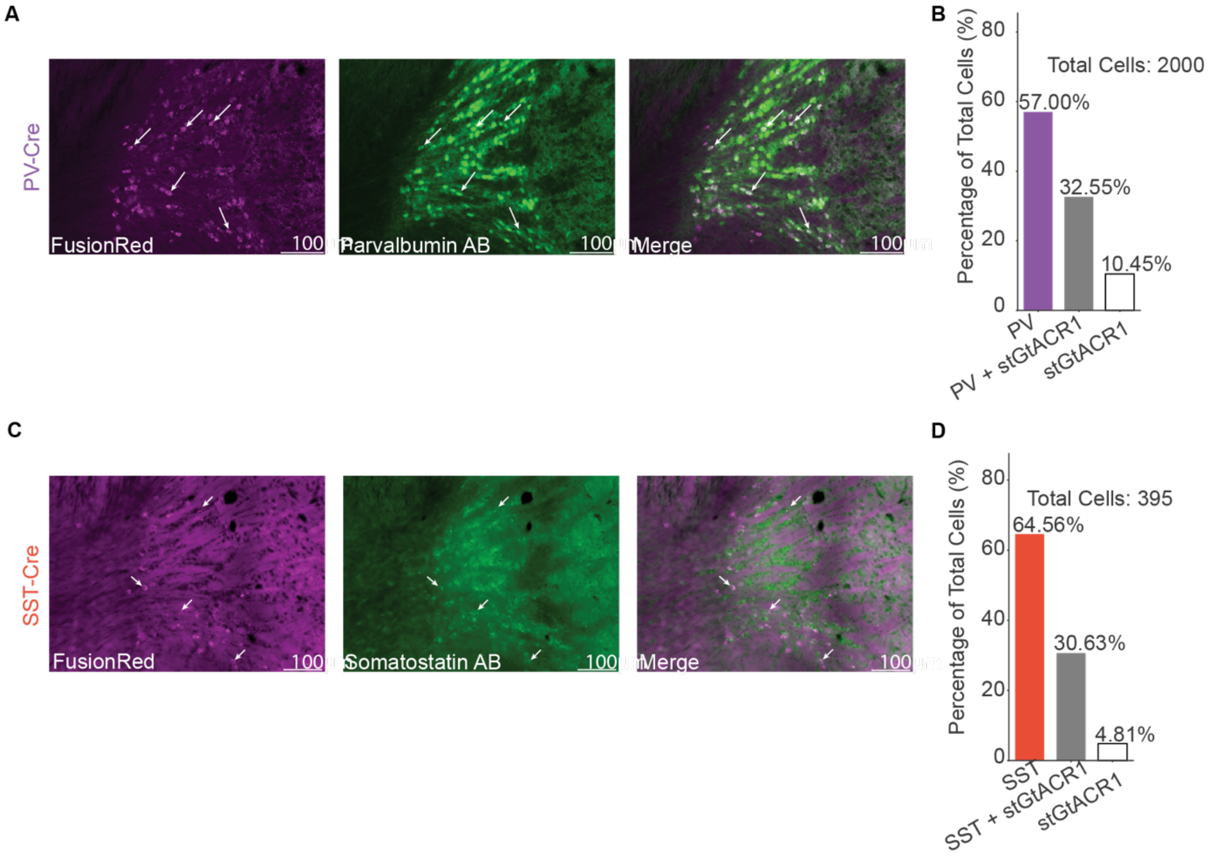


**Supplementary Figure S1. Co-expression of PV+ and SST+ neurons with the inhibitory soma targeted opsin stGtACR1.FusionRed. A.** *Left*: AAV5.hSyn.SIO.stGtACR1.FusionRed expression in the audTRN of a PV-Cre mouse. *Middle*: Immunohistochemical expression of parvalbumin-positive neurons. *Right*: Merged image of PV-audTRN neurons expressing both the parvalbumin antibody and the inhibitory opsin. **B**. Quantification of PV+ neurons, PV+stGtACR1.FusionRed+ neurons, and stGtACR1.FusionRed+ neurons. **C.** *Left*: AAV5.hSyn.SIO.stGtACR1.FusionRed expression in the audTRN of an SST-Cre mouse. *Middle*: Immunohistochemical expression of somatostatin-positive neurons. *Right*: Merged image of SST-audTRN neurons expressing both the somatostatin antibody and the inhibitory opsin. **D**. Quantification of SST+ neurons, SST+stGtACR1.FusionRed+ neurons, and stGtACR1.FusionRed+ neurons. *Scale bar = 100µm. Arrows represent example cells. N=2 mice for each group.*

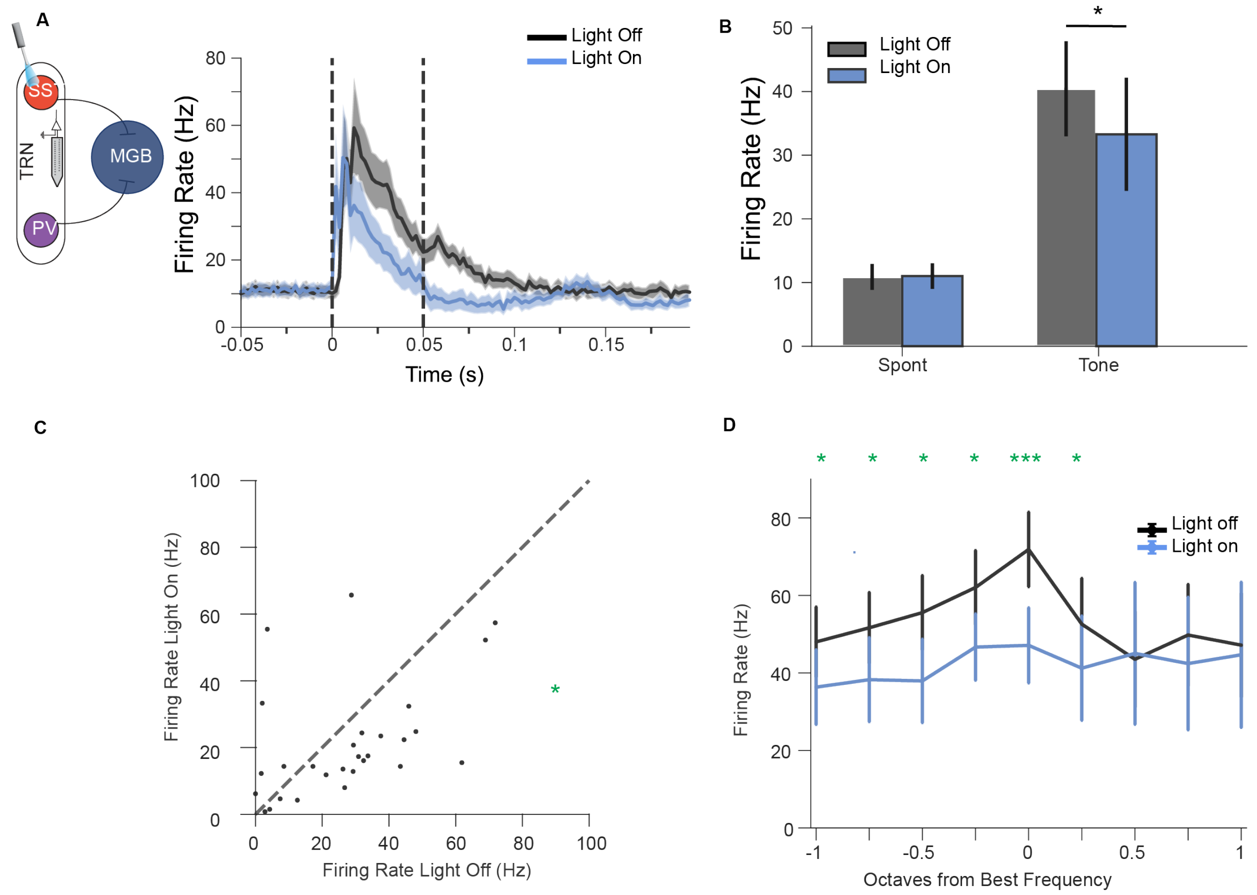


**Supplementary Figure S2. Optical inactivation of SST$^{TRN}$ neurons suppresses tone-evoked activity in the TRN. A.** Average firing rate of recorded units in the TRN in response to 50 ms tone, with vs without optical inactivation of SST neurons (AAV5-hSyn1-SIO-stGtACR1-FusionRed in SST-cre mouse). n= 27 units. Shaded areas represent SEM±. **B.** Average firing rates of TRN units during baseline (Spont), tone-evoked responses (Tone) and offset responses (Offset), with vs without optical inactivation of SST neurons. ***p<0.001, Wilcoxon signed-rank tests. Error bars represent SEM±. **C.** Average firing rate (FR) of TRN units in response to tones on light ON versus light Off trials (with and without optical inactivation of SST neurons). ***p<0.001, Wilcoxon signed-rank test. **E.** Average frequency response function of TRN units with vs without optical inactivation of SST neurons. ***p<0.001, **p<0.01, *p<0.05, Wilcoxon signed-rank tests.

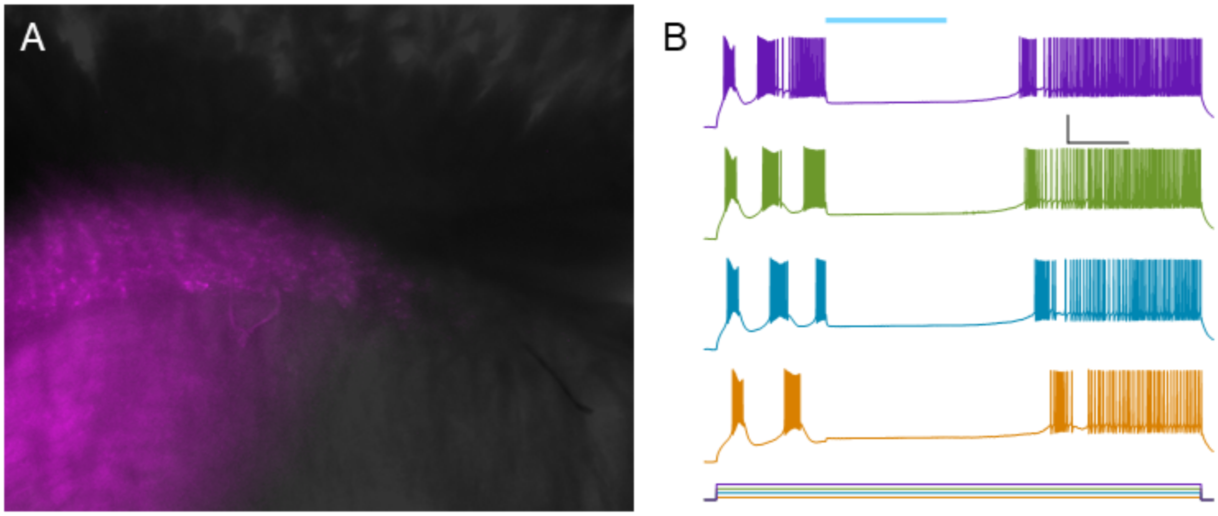


**Supplementary Figure S3. A.** Overlay of live IR and fluorescence images of slice from PV-Cre mouse injected with AAV5_hSyn1-SIO-stGtACR1-FusionRed. **B.** Whole-cell recordings from a TRN neuron from (A) expressing GtACR1, injected with varying amplitudes of current (bottom) and exposed to widefield LED stimulation (blue bar). $V_m$ = -80 mV. Scale bar 25 mV, 250 ms.

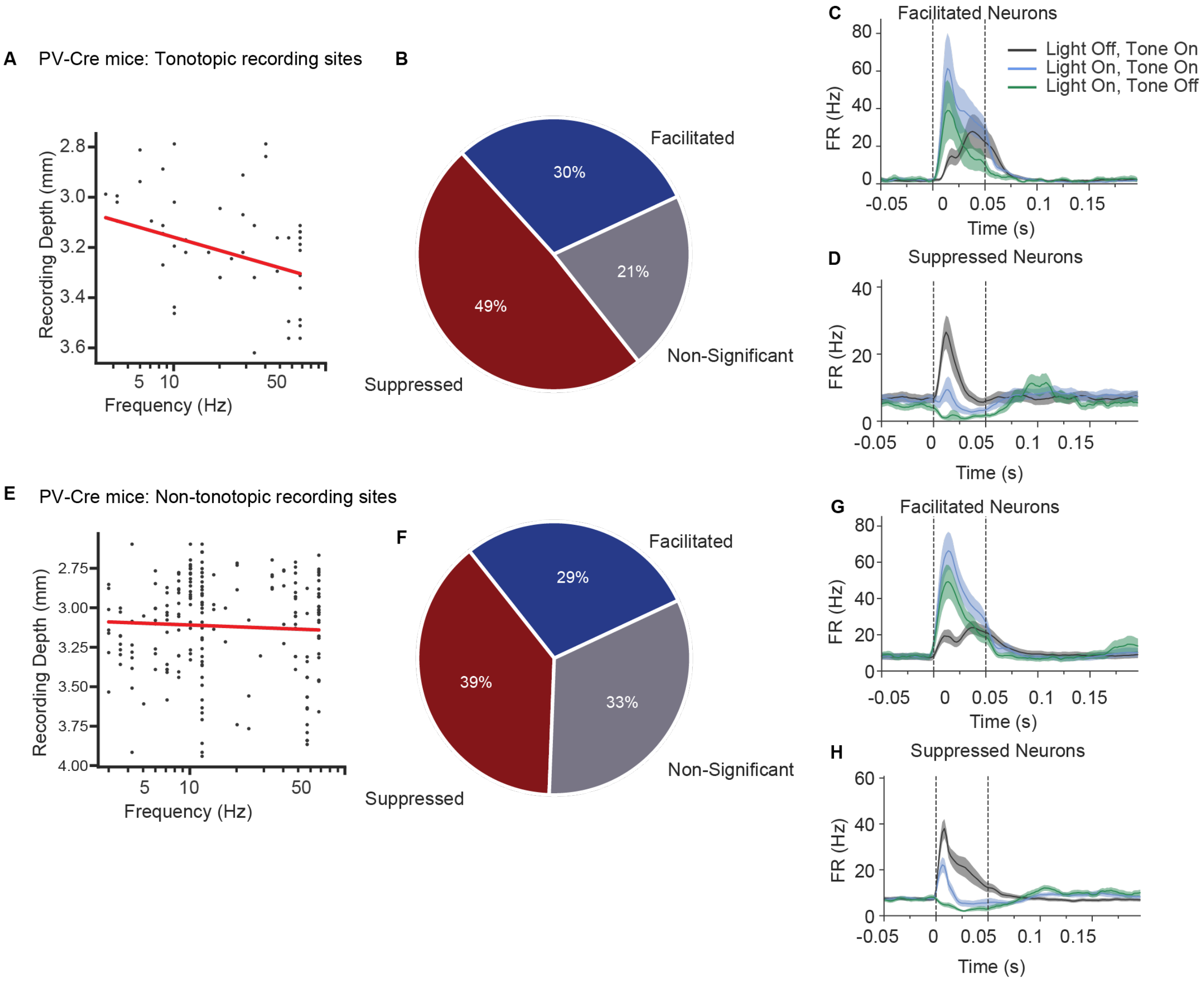


**Supplementary Figure S4. Similar tone and optogenetic responses for recorded MGB neurons in tonotopic and non-tonotopic recording sites in PV-cre mice.** A. Plot of neurons' best frequency vs. recording depth for tonotopic recording sites; N= 47 neurons, 2 recording sites, 2 mice. Red line shows linear regression; r = .41, p = .0047. **B.** Pie chart breaking down the effect of $PV^{TRN}$ inactivation in tonotopic recordings. N= 14 suppressed neurons (49%), 23 facilitated neurons (30%), 10 non-significant neurons (21%). **C, D.** Mean PSTH of facilitated (C) and suppressed (D) recorded neurons in MGB. Light-only trials (green line), tone-only trials (black line), and tone- and laser-on trials (light blue line). Shaded areas represent SEM±. **E-H.** Same as A-F, but for non-tonotopic recording sites. **G.** N= 209 neurons, 10 recording sites, 3 mice; r=-0.06, p=0.39. **H.** N= 81 suppressed neurons (39%), 60 facilitated neurons (29%), 68 non-significant neurons (33%).

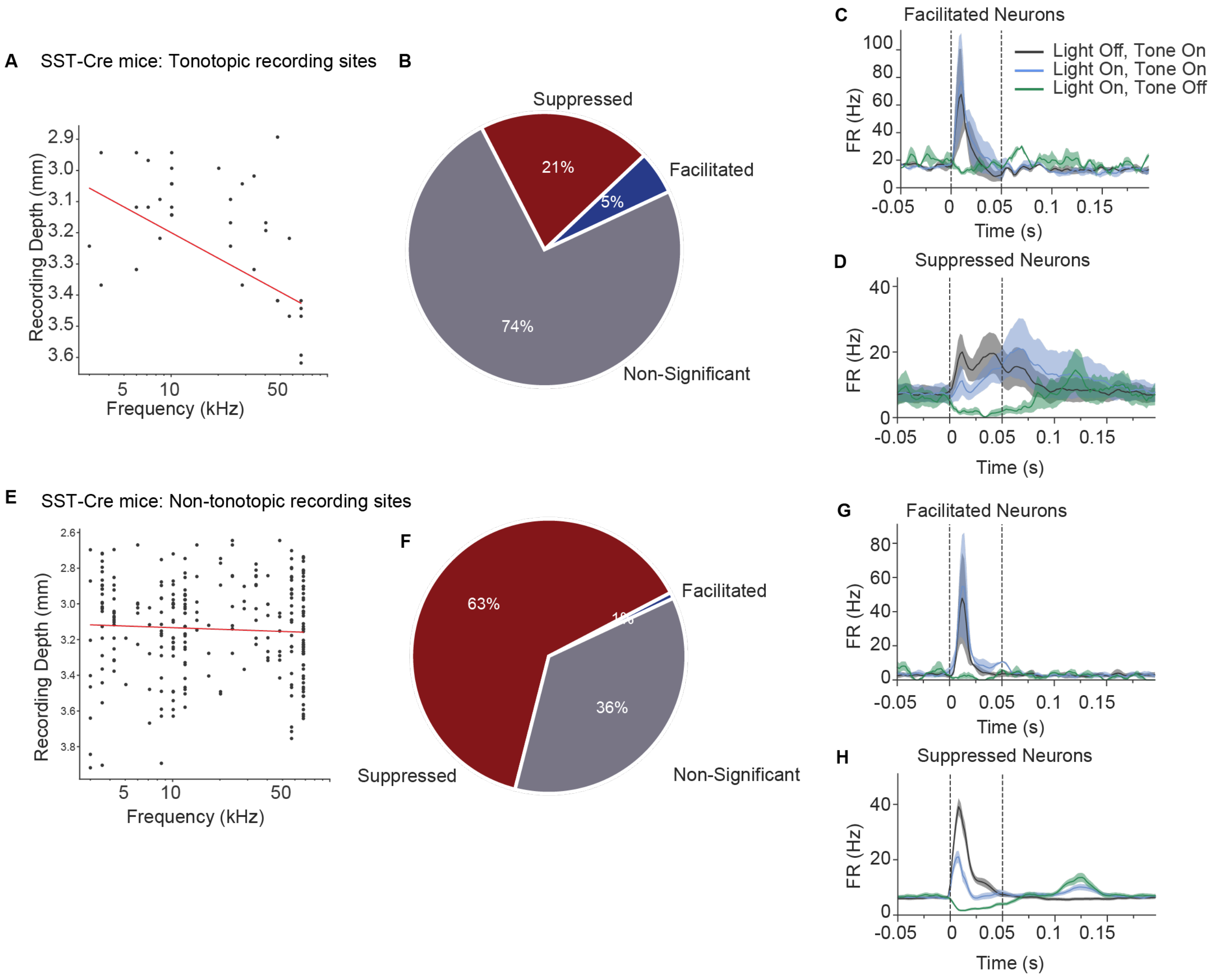


**Supplementary Figure S5. Greater proportion of light-modulated MGB neurons in non-tonotopic than tonotopic recording sites during SST inactivation.** A. Plot of neurons' best frequency vs recording depth for tonotopic recording sites; N= 39 neurons, 1 recording sites, 1 mouse. Red line shows linear regression; r = .67, p = 3e-6. **B.** Pie chart breaking down the effect of $SST^{TRN}$ inactivation in tonotopic recordings. N= 2 facilitated neurons (5%), 8 suppressed neurons (21%), 29 non-significant neurons (74%). **C,D.** Mean PSTH of facilitated (E) and suppressed (F) recorded neurons in MGB. Light-only trials (green line), tone-only trials (black line), and tone- and laser-on trials (light blue line). Shaded areas represent SEM±. **E-H.** Same as A-F, but for non-tonotopic recording sites. **G.** N= 273 neurons, 9 recording sites, 4 mice; r=0.062, p=0.31. **H.** N= 173 suppressed neurons (63%), 2 facilitated neurons (1%), 98 non-significant neurons (36%).

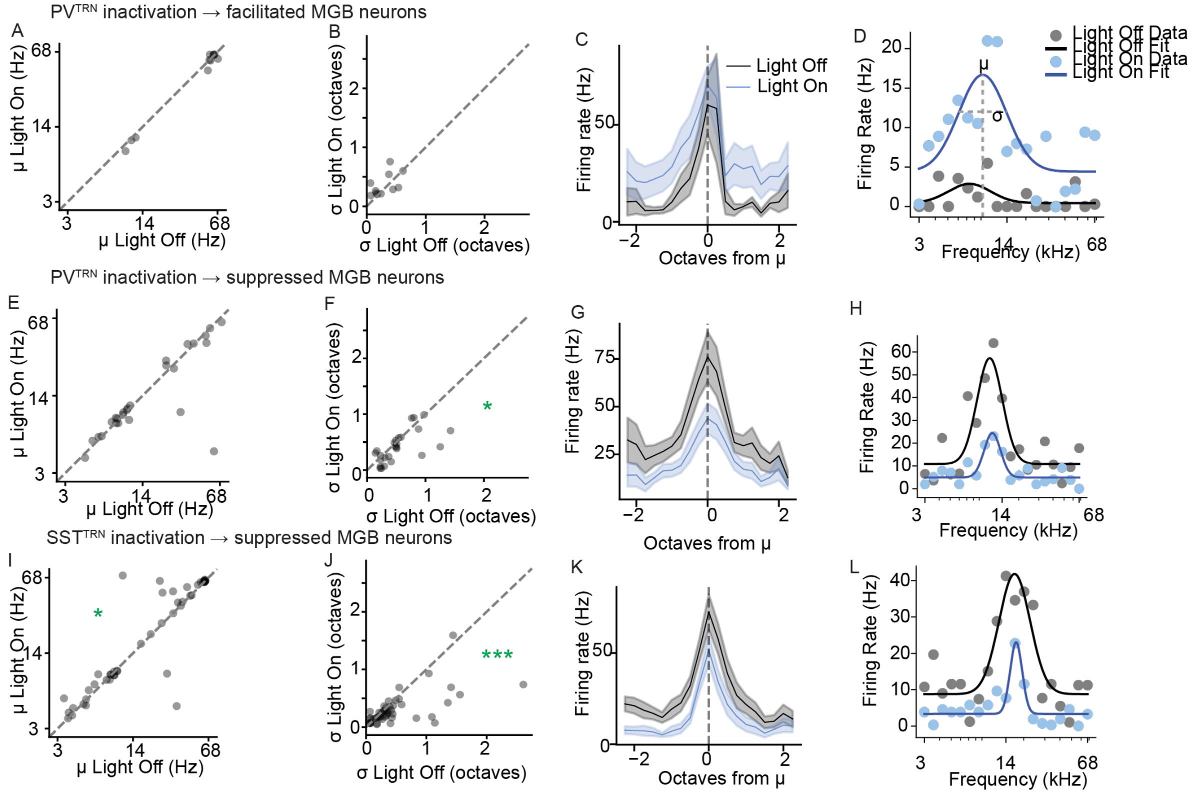


**Supplementary Figure S6. Inactivation of $PV^{TRN}$ and $SST^{TRN}$ neurons shapes frequency tuning in MGB neurons. A.** scatterplot of μ for light-On and light-Off conditions for neurons facilitated by $PV^{TRN}$ inactivation (first row) and neurons suppressed by $PV^{TRN}$ inactivation (second row). **B.** Scatterplots of σ for light-On and light-Off conditions. **C.** Mean frequency response function for light-On (light blue) and light-Off (light gray) trials for neurons centered on μ (0 octaves = μ). **D.** A sample tuning curve for light-On (light blue) and light-Off (grey) conditions along with the gaussian fit for each curve. E-H. Sams as A-D, for suppressed MGB neurons during $PV^{TRN}$ inactivation. **I-L.** Same as *A-D* for $SST^{TRN}$ inactivation, suppressed MGB neurons. Facilitated neurons not show because only N = 1 neuronal responses were well-fitted by a Gaussian for both light on and light off conditions.

We first evaluated how well each tuning curve fit a Gaussian function in both the light-On and light-Off conditions (see Methods). We found that overall the number of MGB neurons with good Gaussian fits ($R^2 > 0.6$ and μ/σ within tested range) was greater in the light-Off condition (n = 33/74 facilitated units and 58/104 suppressed units during $PV^{TRN}$ inactivation; n = 84/181 suppressed units during $SST^{TRN}$ inactivation) compared to the light-On condition (19/74 facilitated units and 43/104 suppressed units during $PV^{TRN}$ inactivation; n = 88/181 suppressed neurons during $SST^{TRN}$ inactivation) in all groups, except for neurons facilitated by $SST^{TRN}$ activation (light-Off, n = 1/4 neurons, light-On, n = 1/4 neurons).

We found no differences in μ (best frequency) for neurons that had good fits in both light-On and light-Off conditions. We did not find significant differences between groups (units facilitated during $PV^{TRN}$ inactivation: N = 11, t = 0.94, p = 0.37; units suppressed during $PV^{TRN}$ inactivation: N = 26, t = 1.36, p = 0.186; units suppressed during $SST^{TRN}$ inactivation: N = 47, t = -1.47, p = 0.147). MGB neurons that were suppressed by $PV^{TRN}$ inactivation showed a significant decrease in σ (PV: N = 26, t = 2.42, p = 0.027; SST: N = 47, t = 3.30, p = 0.0002). However, for neurons facilitated by $PV^{TRN}$ inactivation, there was no change in σ between light-On and light-Off conditions (N = 11, t = -1.27, p = 0.278) and there were no well-fit facilitated neurons during $SST^{TRN}$ inactivation.

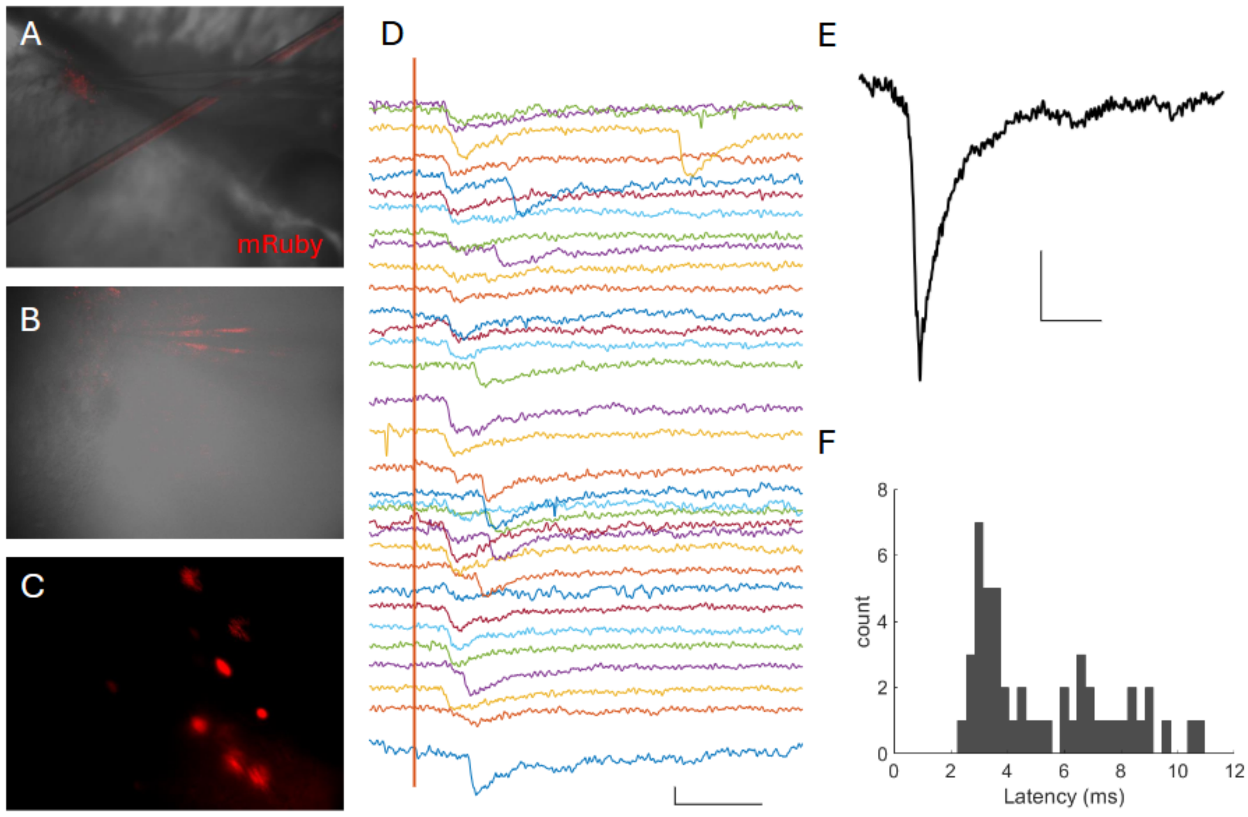


**Supplementary Figure S7. Optogenetic identification of TRN$^{SST}$ to TRN$^{nonSST}$ inhibitory synapse. A.** Widefield image of brain slice containing TRN from an SST-Cre mouse injected with AAV9-DIO-stChrome-mRuby. **B.** 40x image from A with patched neuron; no mRuby expression in this field. **C.** mRuby expression in another field from A confirming neuronal expression. **D.** Inward responses (IPSCs) from neuron patched with high-$Cl^-$ internal solution following widefield optogenetic excitation (red bar). Scale bar 30 pA, 10 ms. Traces are offset for clarity. **E.** Peak-aligned average of responses from D. Scale bar 10 pA, 10 ms. Decay of the IPSC was best fit by a dual exponential with time constants 3.9 and 275 ms. **F.** Latencies of IPSCs from D.

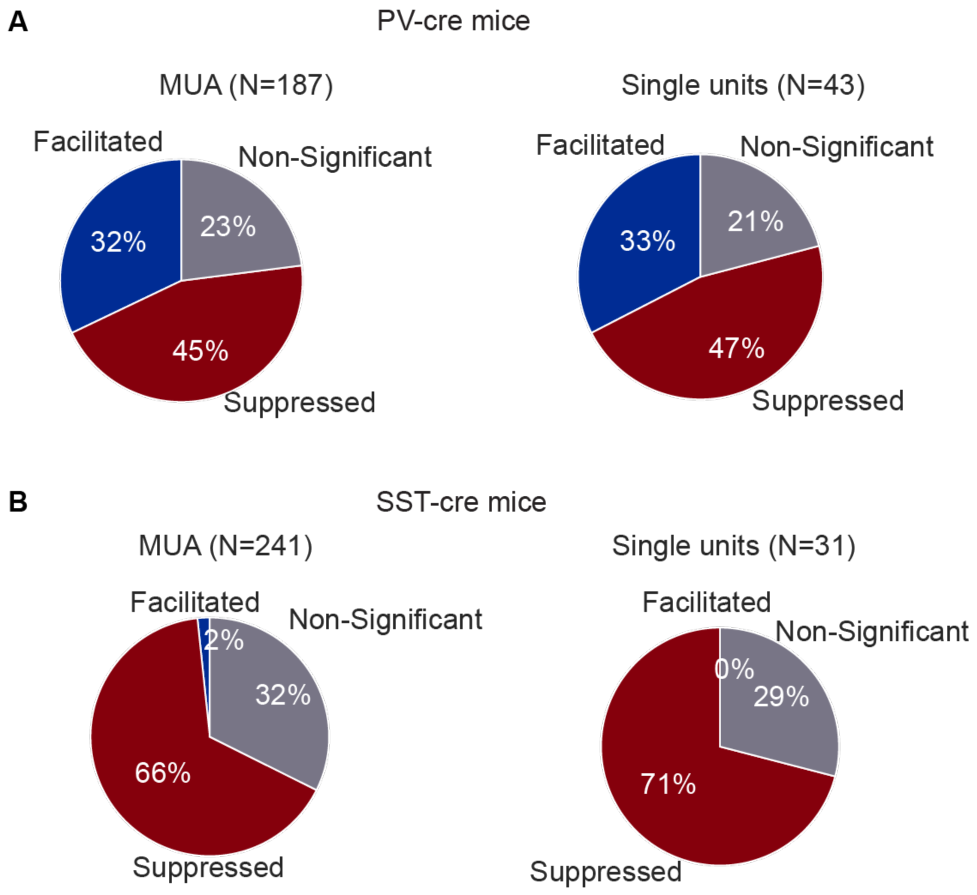


**Supplementary Figure S8. Proportion of facilitated and suppressed MGB single units and multi-units. A.** Proportion of MGB MUA clusters (left) and single units (right) that were facilitated, suppressed, and non-significantly modulated by $PV^{TRN}$ inactivation. **B**. Proportion of MGB MUA clusters (left) and single units (right) that were facilitated, suppressed, and non-significantly modulated by $SST^{TRN}$ inactivation.

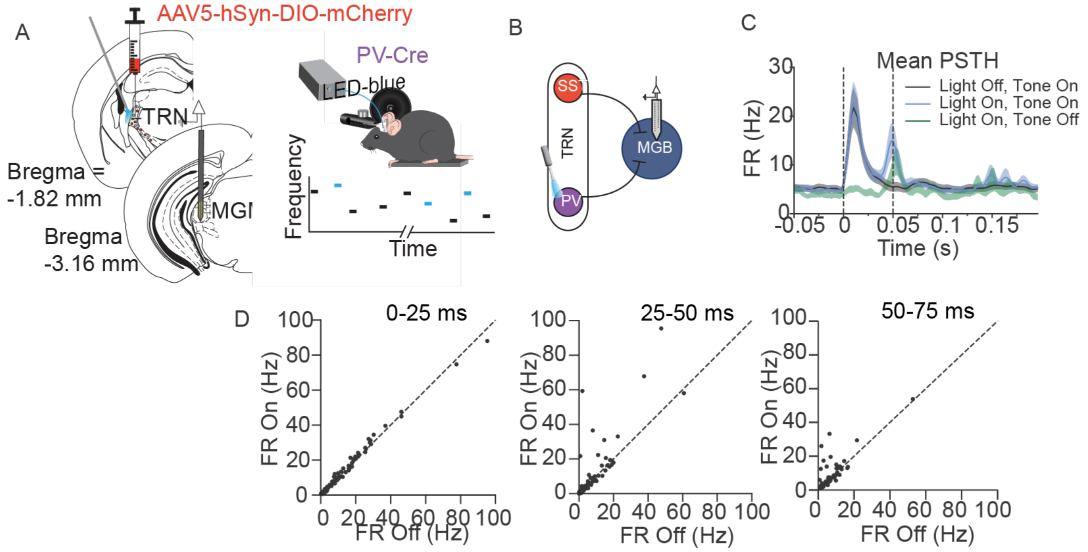


**Supplementary Figure S9. Control experiments for optogenetic manipulation of $PV^{TRN}$ neurons. A.** PV mice were injected with an AAV5-hSyn1-DIO-mCherry virus into the audTRN and implanted with an optic fiber. We recorded from MGB neurons in awake head-fixed mice. We presented a set of pure tones in light-on and light-off conditions. **B**. We presented light in the audTRN of PV mice while recording from neurons in the MGB using a vertical multi-channel electrode that spanned the depth of dMGB and vMGB. **C.** Mean PSTH of recorded neurons in MGB. Light-only trials (green line), tone-only trials (black line), and tone- and laser-on trials (light blue line). **D**. Scatter plot of the mean FR for laser off and laser on conditions 0-25 ms (left), 25-50 ms (center), 50-75 ms (right) post-stimulus onset. *Shaded areas represent SEM±. N=85, n = 2; n.s. for all periods.*

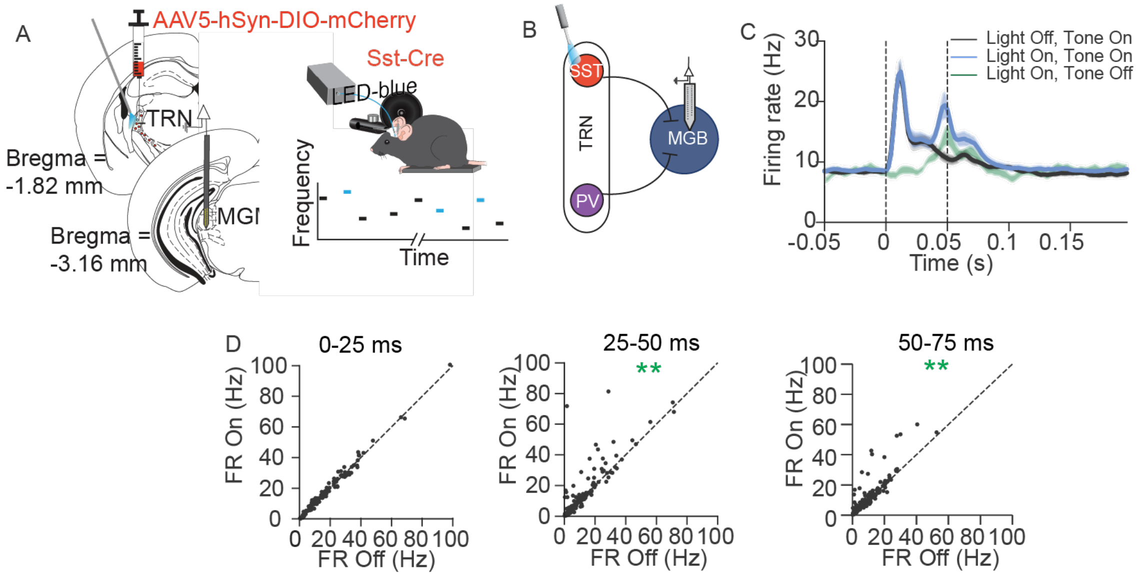


**Supplementary Figure S10. Control experiments for optogenetic manipulation of $SST^{TRN}$ neurons. A.** SST mice were injected with an AAV5-hSyn1-DIO-mCherry virus into the audTRN and implanted with an optic fiber. We recorded from MGB neurons in awake head-fixed mice. We presented a set of pure tones in light-on and light-off conditions. **B**. We presented light in the audTRN of SST mice while recording from neurons in the MGB using a vertical multi-channel electrode that spanned the depth of dMGB and vMGB. **C.** Mean PSTH of recorded neurons in MGB. Light-only trials (green line), tone-only trials (black line), and tone- and laser-on trials (light blue line). **D**. Scatter plot of the mean FR for laser off and laser on conditions 0-25 ms (left), 25-50 ms (center), 50-75 ms (right) post-stimulus onset. *Shaded areas represent SEM±. N=137, n = 2; p-value = n.s. (0-25 ms); 0.006 (25-50 ms), p = 0.000002 (50-75 ms).*
